# 1-Deoxysphingolipids dysregulate membrane properties and cargo trafficking in the early secretory pathway

**DOI:** 10.1101/2025.05.13.652513

**Authors:** Yi-Ting Tsai, Nicolas-Frédéric Lipp, Olivia Seidel, Riya Varma, Aurelie Laguerre, Kristina Solorio-Kirpichyan, Adrian Wong, Roberto J. Brea, Grace H. McGregor, Thekla Cordes, Neal K. Devaraj, Lars Kuerschner, Sonya Neal, Christian M. Metallo, Itay Budin

## Abstract

1-Deoxysphingolipids are non-canonical sphingolipids linked to several diseases, but their cellular effects are poorly understood. Here, we utilize lipid chemical biology approaches to investigate the role of 1-deoxysphingolipid metabolism on the properties and functions of secretory membranes. We first applied organelle-specific bioorthogonal labeling to visualize the subcellular distribution of metabolically tagged 1-deoxysphingolipids in RPE-1 cells, observing that they are retained in the endoplasmic reticulum (ER). We found that 1-deoxysphingolipids can be transported by the non-vesicular transporter CERT *in vitro* but are retained at ER exit sites (ERES) in cells, suggesting that they do not efficiently sort into vesicular carriers. Cells expressing disease-associated variants of serine palmitoyl-CoA transferase (SPT) accumulated long-chain 1-deoxysphingolipids, which reduced ER membrane fluidity and enlarged ERES. We observed that the rates of membrane protein release from the ER were altered in response to mutant SPT expression, in a manner that was dependent on the cargo affinity for ordered or disordered membranes. We propose that dysregulation of sphingolipid metabolism alters secretory membrane properties, which can then modulate protein trafficking.

## Introduction

Sphingolipids (SLs) represent a major class of cell membrane and signaling lipids whose dysregulation is associated with a range of metabolic diseases. SL biosynthesis is initiated by the condensation of L-serine with fatty acyl-CoAs by Serine Palmitoyl-CoA Transferase (SPT), an heteromeric ER membrane protein complex consisting of both catalytic and regulatory subunits^1^. The ketosphinganine product of SPT is rapidly reduced to sphinganine (SA), the building block of all SLs (**Figure 1A**). From its L-serine precursor, the sphingoid base of SA retains a secondary amine, which is *N*-acylated to form dihydroceramides (DHCers) that are *trans* desaturated at the 4,5 position into ceramides, and a primary hydroxyl group at the C1 position, which is modified by polar groups at the Golgi apparatus^2^ to form the complex SLs sphingomyelin (SM) and glycosphingolipids (glycoSL), like glucosyl ceramide (GlcCer), galactosyl ceramide (GalCer), lactosyl ceramides (LactosylCer), and gangliosides. Complex SLs are enriched in the exoplasmic/luminal leaflet of late secretory vesicles and contribute substantially to the ordered-nature of these bilayers^3^.

**Figure 1.**
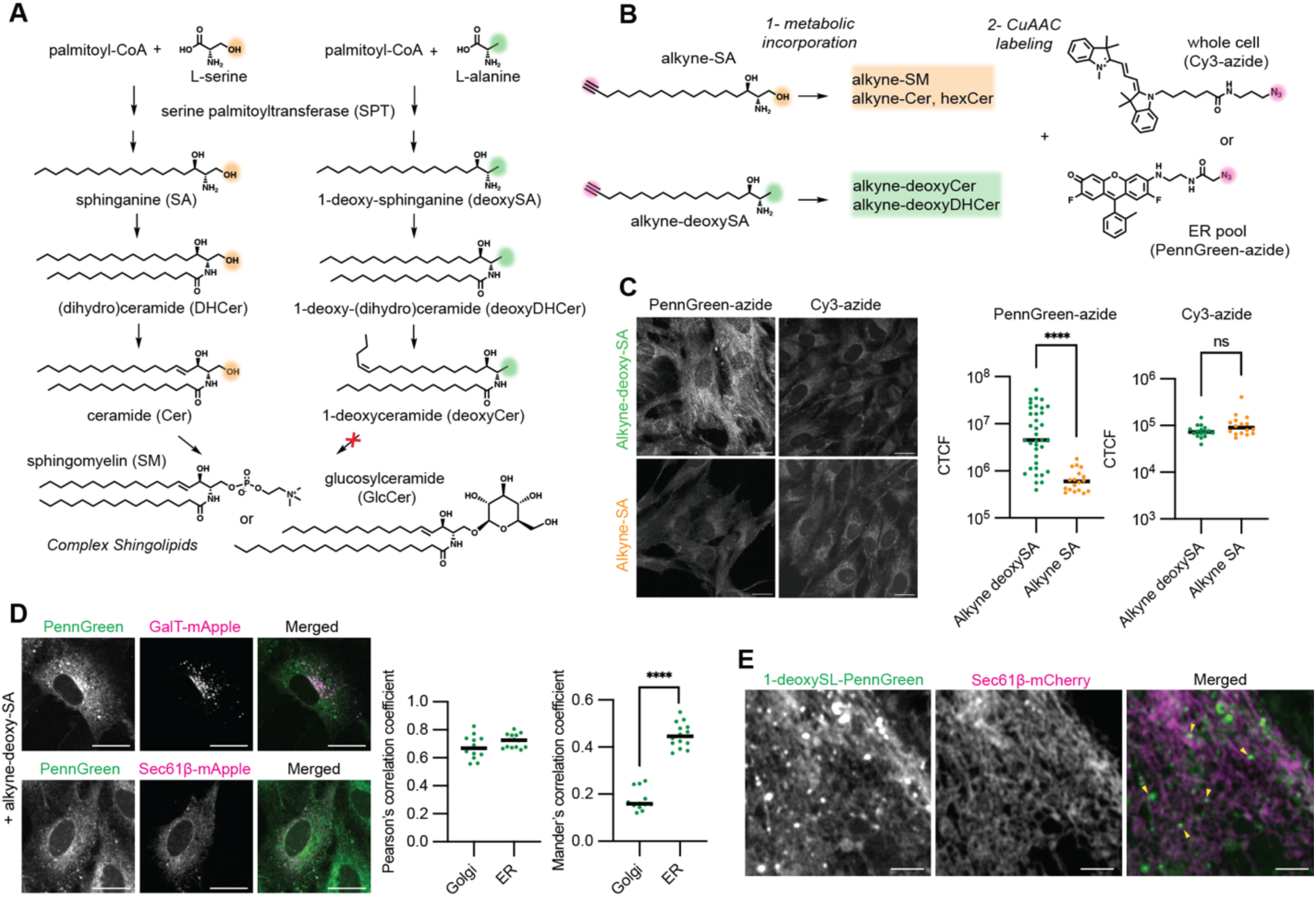
Organelle-specific labeling of 1-deoxySLs identifies their accumulation in early secretory membranes. **A**. Schematic of SL and 1-deoxySL biosynthetic pathway depicting the intermediate lipid species and their chemical structures. **B**. Schematic of the Cu(I)-catalyzed azide-alkyne cycloaddition (CuAAC) approach utilized in this study to identify the localization of 1-deoxySL in subcellular compartments. RPE-1 cells were fed either with either 0.1 μM alkyne-deoxySA, a precursor for 1-deoxySL biosynthesis with an alkyne handle, or 0.1 μM alkyne-SA, and incubated for 17 hours to facilitate the metabolic incorporation, after which cells were fixed and CuAAC reaction was carried with either PennGreen-azide, for specific localization of the lipid in early secretory membranes or Cy3-azide, for labeling of the alkyne lipid in all cell membranes. To compare with the distribution of canonical SL products, alkyne-SA labeling was performed in parallel. **C.** Imaging of RPE-1 cells fed with either alkyne-deoxySA or alkyne-SA and reacted with PennGreen-azide or Cy3-azide. Scale bar, 20 μm. Alkyne-deoxySA treatment led to a larger PennGreen-azide corrected total cell fluorescence (CTCF) after washing of unreacted dye, indicating an accumulation of 1-deoxySL in the early secretory pathway. Each point represents an individual field of cells acquired and processed identically across two biological replicates. ****, P < 0.0001 by Mann-Whitney test. In contrast, the reaction with Cy3-azide did not show increased staining for the alkyne-deoxySA. **D.** Colocalization analysis of 1-deoxySL-PennGreen products with Golgi (GalT-mApple) and ER (Sec61β-mApple) markers. Represent images are shown to the left and quantification of the Pearson’s and Mander’s correlation coefficients to the right. PennGreen fluorescence intensity is strongly correlated with both markers, but overalaps more with the ER. Each point represents an individual cell. ****, P < 0.0001 by Mann-Whitney test. Scale bar, 20 μm. **E.** Airyscan images showing that co-localization of 1-deoxySL-PennGreen products with the ER network labeled by Sec61β-mCherry, with additional punctae that are not labeled by Sec61β-mCherry dispersed through the network indicated by the arrows in the merged image. Scale bar, 2.5 μm.

SPT can also incorporate lower-affinity alternative amino acids in place of L-serine, most commonly L-alanine. The resulting products – 1-deoxy ketosphinganine and 1-deoxy sphinganine (deoxySA) – lack the C1-hydroxyl on the sphingoid base. They are *N*-acylated to form 1-deoxydihydroceramide (deoxyDHCer), which is desaturated into 1-deoxyceramide (deoxyCer), although the resulting double bond is likely *cis* and positioned further down the sphingoid chain than the Cer^4^. Unlike canonical SLs, dihydro species accumulate in the 1-deoxy pathway, likely reflecting C1 hydroxyl substrate specificity for dihydroceramide desaturase (DEGS1). These 1-deoxysphingolipids (1-deoxySLs) also cannot be further modified into complex SLs, like sphingomyelin (SM) or glycosphingolipids, nor broken down by the canonical SL degradation pathway^5^, both of which require modifications on the C1 hydroxyl group. They thus represent an alternative branch of SL biosynthesis^6^. 1-DeoxySLs, especially the dihydro species (deoxDHCer) that are abundant in cells, retain the high melting temperature of ceramides but are even less miscible due to their extreme hydrophobicities^7,8^.

While first identified in mollusks^9^ and pathogenic fungi^10^, 1-deoxySLs have more recently been recognized as endogenous metabolites in mammalian cells and as potential drivers of human disease^11^. In mammals, 1-deoxySL metabolism is controlled by amino acid concentrations and the substrate selectivity of SPT. Depletion of serum L-serine levels causes accumulation in patients with type 2 diabetes mellitus (T2DB), for which 1-deoxySLs serve as a biomarkers^12^ and may function in associated diabetic sensory polyneuropathy (DSN)^13^. Specific alleles of SPT subunits that lose substrate selectivity for L-serine can also drive disease metabolism. Mutations in the SPT catalytic subunits SPTLC1 or SPTLC2 are causative for hereditary sensory and autonomic neuropathy type 1 (HSAN1), an axonal neuropathy whose severity is correlated with plasma 1-deoxySL levels^14^. When expressed in cells, HSAN1-associated alleles in SPT subunits cause large increases in substrate affinity for L-alanine vs. L-serine^15^. Expression of SPTLC1 disease-variants was shown to drive pathologies in animal models^16^, which are rescued by heterozygous overexpression of wild-type alleles^17^. Patients with HSAN1 also show high susceptibility to develop macular telangiectasia type 2 (MacTel)^18^, which manifests in adult-onset blindness and correlates with low serum L-serine levels^19^. In MacTel patients, the macula features distinct regions of pigment loss, while the subretinal space between photoreceptors and the Retinal Pigment Epithelium (RPE) accumulates rod outer segments (ROS) and other cell debris^20^. Such subretinal deposits are also signatures of other forms of macular degeneration^21^. MacTel patient RPE cells show reduced cell surface expression of MerTK^22^, an apical membrane receptor for phagocytic recycling of ROS^23^, suggesting a potential mechanism for the accumulation of subretinal debris.

Mechanisms underlying 1-deoxySL toxicity in cells or tissue-specific phenotypes in patients are still unresolved. Phenotypic effects of 1-deoxySLs are most commonly assayed in cell culture by medium supplementation with exogenous 1-deoxySA, comparing its effects with equivalent concentrations of SA. After uptake, 1-deoxySA is converted into the corresponding 1-deoxyDHCer and 1-deoxyCers that localize to a range of cellular compartments, including mitochondria, Golgi, and lysosomes, as imaged by bioorthogonal reactive handles^24^. Resulting toxicities vary by cell type, with cancer cells^25^ and neurons^26^ being particularly sensitive. Reported molecular phenotypes upon supplementation with μM-concentrations of 1-deoxySA include the induction of the unfolded protein response (UPR)^27^, pro-apoptotic PKC signaling^25,28^, autophagosome formation^29^, and loss of mitochondrial respiration^5^. Supplementation of 1-deoxySA depends on cell permeability for single-chain SLs, which are expected to diffuse between intracellular membranes^30^. In contrast, endogenous SA is produced in the ER and then metabolized to insoluble double-chained SL products, all of which must be trafficked by lipid transport pathways. 1-DeoxySL accumulation is higher in typical 1-deoxySA supplementation experiments than from expression of disease-associated SPT variants^31^. Thus, it is not clear if exogenously added SL metabolites fully reflect the localization and phenotypes of those that are endogenously produced. An alternative approach to investigating 1-deoxySL function is the study of patient-derived cell lines with SPT mutations^32^ or heterologous expression of these SPT variants in engineered cell lines^31^.

Here we apply a combination of chemical and cell biology approaches to investigate the role of 1-deoxySLs in the secretory pathway of cells. We were motivated by the reduced surface display of membrane proteins in the RPE of MacTel patients to consider the role of 1-deoxySLs metabolism in secretory pathway function. Utilizing an immortalized RPE cell line as a model, we report that 1-deoxySLs accumulate in the early secretory pathway, where they alter ER membrane fluidity and protein cargo export.

## Results

### 1-DeoxySLs accumulate in the early secretory pathway

The most abundant 1-deoxySLs, deoxyCers and deoxyDHCers, are synthesized in the ER alongside their corresponding SL precursors. Ceramides are efficiently exported from the ER membrane by both vesicular and non-vesicular pathways^2^ and we asked if this was the case for their 1-deoxy counterparts. Bioorthogonal ‘click’ chemistry is one approach to image subcellular distributions of specific membrane lipids, provided that tracers can be fed to cells that can be readily incorporated into native metabolic pathways and feature minimally-disruptive chemical handles^33^. The single-chain SL precursor SA modified with a terminal alkyne (alkyne-SA), alongside its 1-deoxy counterpart (alkyne-deoxySA) (**Figure 1B**), can be used to visualize SL distributions through Copper(I)-catalyzed Alkyne-Azide Cycloaddition (CuAAC) with azide-containing fluorophores^5^. Previous analyses with alkyne-deoxySA fed cells showed its incorporation into deoxyDHCer and deoxyCer species and staining of several compartments within cells, including the ER, Golgi, mitochondria, and lysosomes^5,29^.

To better visualize the distribution of 1-deoxySLs at their site of synthesis and transport in the early secretory pathway, we sought to employ organelle-targeted bioorthogonal chemistry. The fluorophore Pennsylvania Green has been previously shown to localize predominantly to ER and Golgi membranes through its rhodol group^34^ and used to specifically label azido-labeled lipids^35^ and native proteins^36^ in these compartments. To carry out such an analysis on alkyne-SA and alkyne-deoxySA probes, we synthesized an azido Pennsylvia Green (PennGreen-azide) that is compatible with CuAAC reactions (**Figure S1**). We confirmed that PennGreen-azide localized predominantly to ER and Golgi membranes, with some residual staining of mitochondria as previously observed (**Figure S2A**). Cells were incubated with trace amounts of alkyne-SA and alkyne-deoxySA (0.1 μM) before PennGreen-azide staining, fixation, conjugation, and washing steps to remove non-conjugated fluorophores. Cells not treated with an alkyne lipid showed no residual PennGreen signal after washing steps (**Figure S2B**).

We contrasted labeling patterns after CuAAC reactions of alkyne-SA and alkyne-deoxySA with PennGreen-azide with those utilizing Cy3-azide, a hydrophobic fluorophore that indiscriminately localizes to all membranes within the cell. In all cases, unreactive dyes are washed away, so remaining fluorescent signals reveal the localization of labeled lipid substrates (**Figure 1C**). Canonical SL products, labeled with alkyne-SA, showed robust staining with Cy3-azide, but poor labeling with PennGreen-azide, consistent with their rapid transport from the ER and Golgi to later secretory compartments where they accumulate. In contrast, 1-deoxySLs, labeled with alkyne-deoxySA, showed a strong signal upon labeling with PennGreen-azide, suggesting an enrichment in the early secretory pathway. In alkyne-deoxySA fed cells, the signal for the PennGreen product (PennGreen-1-deoxySL) was correlated with that of both ER (Sec61β) and Golgi (GalT) markers, but predominantly overlapped with the former (**Figure 1D**). These data were broadly consistent with experiments showing that 1-deoxy variants of C6-NBD-Ceramide do not label Golgi compartments and are retained in the ER^37^. Examination of high-resolution Airyscan micrographs showed that PennGreen-1-deoxySL signal colocalized with the peripheral ER network, but also showed accumulation at dispersed punctae that were not labeled with Sec61β (**Figure 1D**).

### 1-deoxySLs accumulation occurs at sites of vesicular trafficking

We next considered the mechanism by which 1-deoxySLs become retained in the ER. Canonical ceramides are trafficked from the ER through vesicular and non-vesicular mechanisms (**Figure 2A)**. The lipid transfer protein (LTP) CERT^38^ localizes to ER-Golgi contact sites^39^ and contributes to SM synthesis by Golgi-localized SMS1^40^. The structure of CERT bound to ceramide ligands shows hydrogen bonding interactions between residues in the (StAR)-related lipid transfer (START) domain and the C1-hydroxyl of ceramide, which is absent in 1-deoxySLs (**Figure 2B)**. To test whether the absence of this interaction alters CERT activity, we synthesized a 1-deoxyDHCer with a C12-NBD *N*-acyl chain (C12-NBD-deoxyDHCer), as well as a version containing the C1 -hydroxyl (C12-NBD-DHCer) for use in FRET-based lipid transfer assays (Materials and Methods). Liposomes containing either C12-NBD-DHCer or C12-NBD-deoxyDHCer as FRET donors and Rhodamine-PE as a FRET acceptor are mixed with non-fluorescent acceptor liposomes. Addition of purified CERT START domain initiated dequenching of the NBD lipids as they were transferred into liposomes lacking Rhodamine-PE (**Figure 2C**). Using this approach, we measured that CERT activity for C12-NBD-deoxyDHCer was ∼2-fold higher than that of C12-NBD-DHCer (**Figure 2D**). Transport rates for both were inhibited by the competitive ceramide analogue HPA-12, indicating that they are trafficked via the same binding pocket. An increased transfer rate upon loss of binding interactions for 1-deoxy ligands is reminiscent of the inverse correlation between ligand affinity and transfer rate observed in Osh/ORP LTPs^41^.

**Figure 2.**
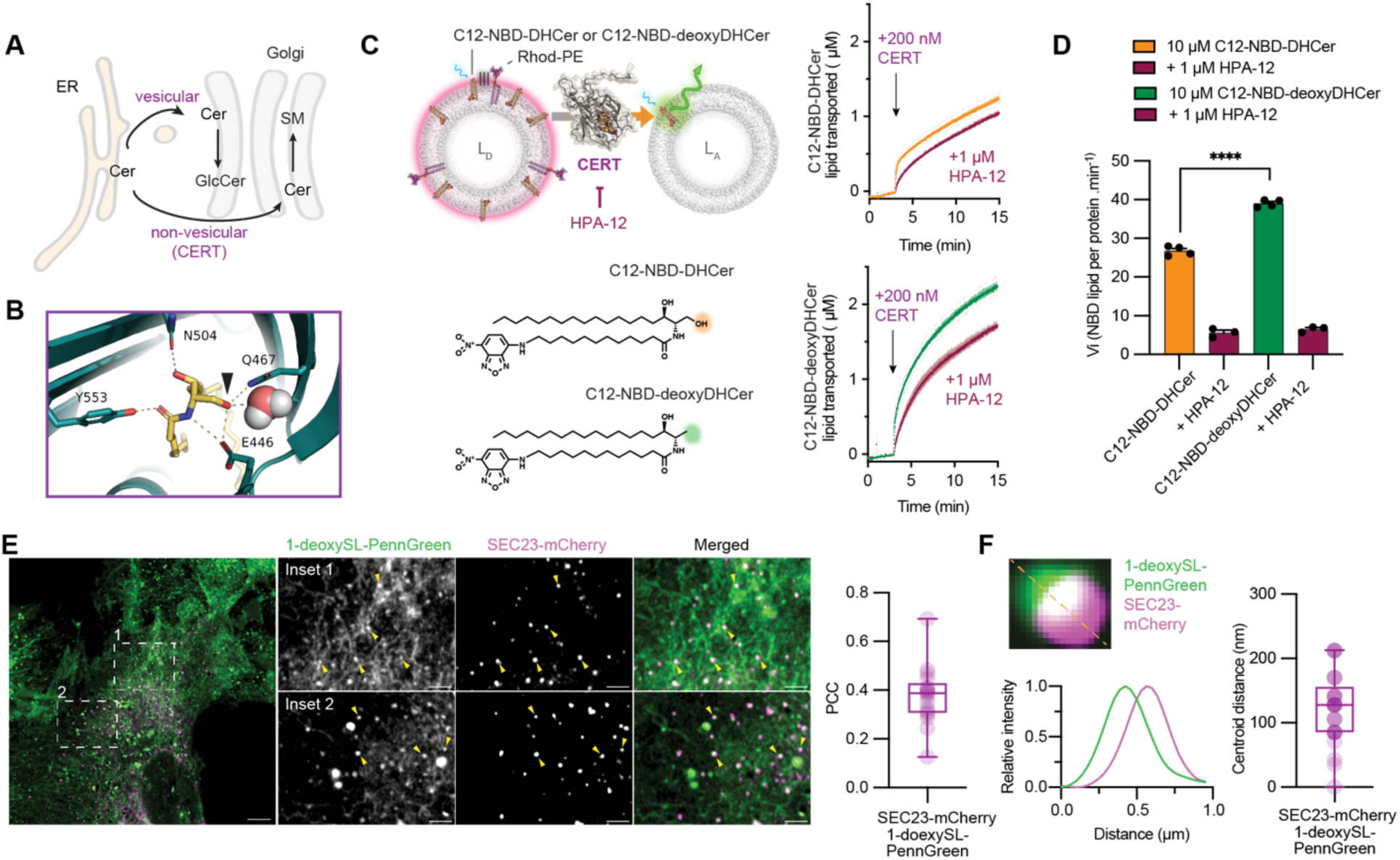
1-DeoxySLs are transported by CERT but accumulate at sites of vesicular trafficking. **A.** Schematic of known Cer trafficking mechanisms in cells. Cer produced in the ER can be exported to the Golgi via vesicular or non-vesicular trafficking, the latter is mediated by the LTP CERT. **B.** The CERT START domain makes polar contacts (Q467, E446, and a coordinated water) with the C1 hydroxyl of Cer (arrow, PDB: 2E3O), which would not be made with 1-deoxy cargoes. **C.** Measurements of CERT transport activity for Cer and 1-deoxyCer cargoes using FRET-based liposome assay (top) and C12-NBD-DHCer/C12-NBD-deoxyDHCer (bottom). Protein preparation and probe synthesis are described in the Materials and Methods. Donor liposomes (L_D_) contained 93 mol% di-oleoyl-phosphatidylcholine (DOPC), 5 mol% of the NBD-labeled lipid, and 2 mol% rhodamine-phosphatidylethanolamine (Rhod-PE). Acceptor liposomes (L_A_) contained only DOPC. **D.** Lipid transport curves for liposomes containing 10 μM C12-NBD-DHCer (top) and C12-NBD-deoxyDHCer (bottom) upon injection of 200 nM CERT to a cuvette containing donor and acceptor liposomes. Absolute lipid transport rates are generated by normalizing FRET donor (NBD) emission with liposome standards. CERT transports C12-NBD-deoxyDHCer faster than C12-NBD-DHCer, and transport of both is slowed in the presence of 1 μM of the competitive inhibition HPA-12. Initial transport rates are reported on the right. Error bars show SEM (N = 4 independent experiments per condition). ****, P < 0.0001 by an unpaired Welch’s t-test. **E.** RPE-1 cells incubated with 0.1 μM 1-deoxySA for 17 hours before fixation, CuAAC labeling with PennGreen-azide, and washing, as in Figure 1. The resulting 1-deoxySL-PennGreen products are observed at punctae that co-localize with the ERES marker SEC23-mCherry. Pearson’s colocalization coefficient (PCC) between 1-deoxySL-PennGreen and SEC23-mCherry is shown for 17 cells. **F.** The centroid of 1-deoxySL-PennGreen punctae overlap with those of SEC23-mCherry, which label ERES, but show an average offset distance of 120 nm. A single example of such overlap is shown with the corresponding two-channel profile intensity along the dashed line. Measurements of centroid-to-centroid distances for 29 exit sites are also shown.

Ceramides are also trafficked from the ER via CERT-independent pathways, which are responsible for glucosylceramide synthesis^42^, and are thought to involve vesicular carriers that bud from the ER membranes at ER exit sites (ERES)^43^. Sorting and trafficking of lipids to ERES is not fully understood but is presumed to rely on their nanometer-scale partitioning into the ERES tubular network^44^. If this segregation is impeded by the reduced solubility and increased hydrophobicity of 1-deoxySLs, we hypothesized that they would accumulate proximal to ERES in cells. The 1-deoxySL-PennGreen punctae we previously observed dispersed through the ER network (**Figure 1E**) co-localized with the COP-II coat protein SEC23, indicating accumulation at or near ERES (**Figure 2E**). High-resolution imaging revealed a randomly oriented offset of 100-200 nm between SEC23-mCherry and 1-deoxySL-PennGreen (**Figure 2F**). This distance is within that measured for the ultrastructure of ERES across a range of cell types^44^. These data suggest that 1-deoxySLs accumulate at regions within or proximal to ERES. A reduced capacity to sort into secretory vesicles or inhibition of their release could thus underlie their retention in the ER.

### A disease-associated SPT variant recapitulates 1-deoxySL overproduction

Compared to other lipid classes, ceramides and, especially, 1-deoxyCer show unusual properties because of their propensity to form high order gel-like membrane domains due to their immiscibility with other lipids *in vitro*^8^, especially for 1-deoxyDH species^7^. If 1-deoxySLs accumulate in the ER, we hypothesize that their overproduction would drive changes to ER structure and function. To promote 1-deoxySL synthesis without altering amino acid availability, which directly induces ER stress^45,46^, we constructed stable cell lines in which 1-deoxySL synthesis is induced through inducible expression of the cysteine 133 to tryptophan allele of SPTLC1 (SPT^C133W^). C133W is the most common mutation identified in HSAN1 patients^47^ and lies at the interface with SPTLC2, near the complex’s PLP cofactor^1^. We expressed SPT^C133W^ using a doxycycline-inducible promoter; as endogenous SPTLC1 was retained, increased 1-deoxySLs production requires SPT^C133W^ overexpression. To control for artifacts involving SPTLC1 expression, we generated an analogous strain harboring the wild-type (WT) variant of SPTLC1 (SPT^WT^) which showed similar expression across a range of doxycycline concentrations (**Figure 3A**). In the absence of doxycycline, growth of both SPT^WT^ and SPT^C133W^ was identical to RPE-1. Only the latter showed a growth defect under induction maximum conditions (**Figure 3B**), albeit a minor one. Thus, SPT overexpression itself is not deleterious to RPE-1 cells under these growth conditions.

**Figure 3.**
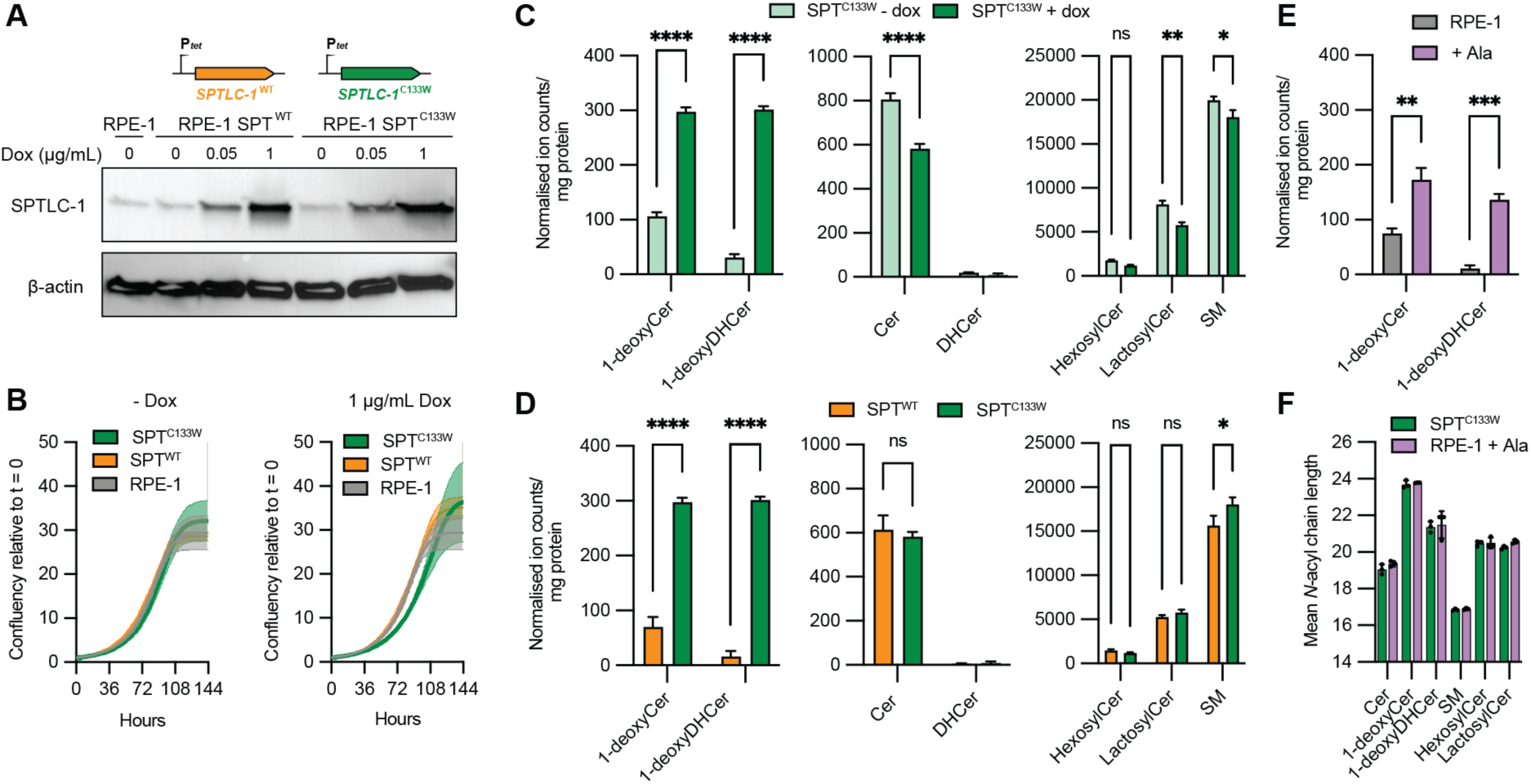
Endogenous overproduction of 1-deoxySLs in RPE-1. **A.** Immunoblotting of SPTLC-1 and β-actin in RPE-1 cell line expressing additional copies of wild type (SPT^WT^, orange) or mutant SPTLC1 (SPT^C133W^, green) under a titratable Tet promoter. Upon incubation with the indicated concentration of doxycycline for 48 hours, cell lysates were obtained and analysed. WT and mutant SPTLC1 show similar expression levels. **B.** Growth of RPE-1, SPT^WT^, and SPT^C133W^ is identical under no induction, while the latter shows a modest growth defect under 1 μg/mL doxycycline induction. Error bars indicate SEM for N = 3 independent wells measured with an IncuCyte system. **C.** Induction of SPT^C133W^ causes high accumulation of 1-deoxy products (left) and depletions of canonical Cers (middle) and complex SLs (right). Levels of canonical SL and deoxySL were compared between SPT^C133W^ cells cultured with or without 1 μg/mL doxycycline for 48h. **D.** Compared to induced SPT^WT^ cells, SPT^C133W^ cells show high accumulation of 1-deoxySLs (left) and only minor changes to canonical Cers (middle) and complex SLs (right). **E.** Moderate accumulation of 1-deoxySLs in RPE-1 cells supplemented with 1 mM L-alanina (Ala) for 48 hours. **F.** In both SPT^C133W^ and Ala-fed RPE-1 cells, 1-deoxyCer and 1-deoxyDHCer species have an average *N*-acyl chain length (mean of 23 carbons) that is longer than canonical Cers (19 carbons). This VLCFA resembles the profile of Hexosyl- and Lactosyl Cers (20 carbons), while SM species (17 carbons) show short *N*-acyl chain lengths. For all lipidomic analyses, error bars indicate SEM (extracts from N = 3 independent culture dishes). *, P < 0.05; ** P < 0.01; ****, P < 0.0001 by 2-way ANOVA.

Lipidomics analyses showed that, under induction, SPT^C133W^ led to an 4 and 10-fold increase in deoxyCers and deoxyDHCers, respectively, over both uninduced SPT^C133W^ cells (**Figure 3C**) and the induced SPT^WT^ control strain (**Figure 3D**). Induced SPT^WT^ and SPT^C133W^ showed identical ceramide and complex sphingolipids lipids, with the exception of a minor increase in SM levels in the latter. In contrast, SPT^C133W^ induction led to moderate decreases in ceramides, lactosyl ceramides, and sphingomyelin, when compared to uninduced cells of the same background (**Figure 3C**). We thus chose the comparison of induced SPT^WT^ and SPT^C133W^, which have highly similar sphingolipid profiles except for 1-deoxyCers and 1-deoxyDHCers levels, for follow up experiments.

In addition to expression of SPTLC1 mutants, 1-deoxySLs can also accumulate in response to changes in the L-alanine (Ala) to L-serine ratio in the medium, mimicking the effect of amino acid dysregulation observed in metabolic disease^48^. In RPE-1, medium supplementation with 1 mM Ala caused accumulation of deoxyCers and deoxyDHCers compared to control RPE-1 (**Figure 3E**). However, deoxyCers and deoxyDHCers levels in Ala-fed RPE-1 were 2-3-fold less than that for induced SPT^C133W^, suggesting that mutations in SPTLC1 can have a larger effect than amino acid concentrations in determining cellular 1-deoxySL levels. In both induced SPT^C133W^ and Ala-supplemented cells, the most abundant 1-deoxySL species were species saturated with 24:0 *N*-acyl chains, e.g. m18:0/24:0 deoxyDHCer and m18:1/24:0 deoxyCer, with other very long chain fatty acid (VLCFA) *N*-acyl chains like 24:1 and 26:0 also abundant (**Figure S3, Figure S4**). This distribution of acyl chains more closely resembled that of glycolipids (hexosyl and lactosyl ceramides), which are also enriched in VLCFAs, than that of SM or canonical Cers, which have shorter *N*-acyl chains (e.g. C16:0) (**Figure 3F**).

In previous studies, high levels of 1-deoxySLs were generated in a variety of models through supplementation with exogenously added 1-deoxySAs at μM concentrations, which caused mitochondrial^24^ and ER stress^27^. We thus tested whether endogenous 1-deoxySL overproduction would cause similar phenotypes. We measured the induction of ER stress response pathways through blotting against canonical Unfolded Protein Response (UPR) markers: activated (phosphorylated) forms of the sensors IRE-1 and Akt, and the ER-associated degradation proteins Derlin-1 and Derlin-3. We did not observe major changes in the abundance of these markers when comparing uninduced and induced SPT^C133W^ and SPT^WT^ cells (**Figure S5A**). We also observed that neither SPT^C133W^ nor SPT^WT^ cells showed reduction in respiratory capacity measured using SeaHorse respirometry, indicating a lack of significant mitochondrial dysfunction (**Figure S5B**). These results indicate that accumulation of 1-deoxySLs in SPT^C133W^ expressing RPE-1 cells is not tied to the mitochondrial and ER stresses previously reported in other systems treated with exogenous 1-deoxySL precursors^5,27^.

### 1-deoxySL accumulation causes altered ER membrane packing and ERES morphology

Given their very different biophysical behaviors compared to bulk phospholipids, we asked if 1-deoxySL could instead alter membrane properties in the ER where they accumulate. In unstressed cells, the ER is largely devoid of high-melting temperature lipids and order-promoting cholesterol due to rapid trafficking of these components after their synthesis^49,50^. We hypothesized that retention of long-chain 1-deoxySLs could thus alter membrane packing in this organelle. To test this, we used the solvatochromic dye Laurdan, whose emission spectrum is blue shifted in more ordered membrane environments. The change in Laurdan emission is quantified by a unit-less Generalized Polarization (GP) ratio based on pixel intensities taken simultaneously at two emission wavelength windows in confocal micrographs^51^. Laurdan labels all lipid environments in the cell, so we used an additional marker (Sec61-mCherry) to auto-mask the ER-specific Laurdan signal (GP_ER_). Using this pipeline, we generated GP_ER_ heatmaps across individual cells and monitored more highly ordered regions by analysis of the emission relative signal across two channels (**Figure 4A**).

**Figure 4.**
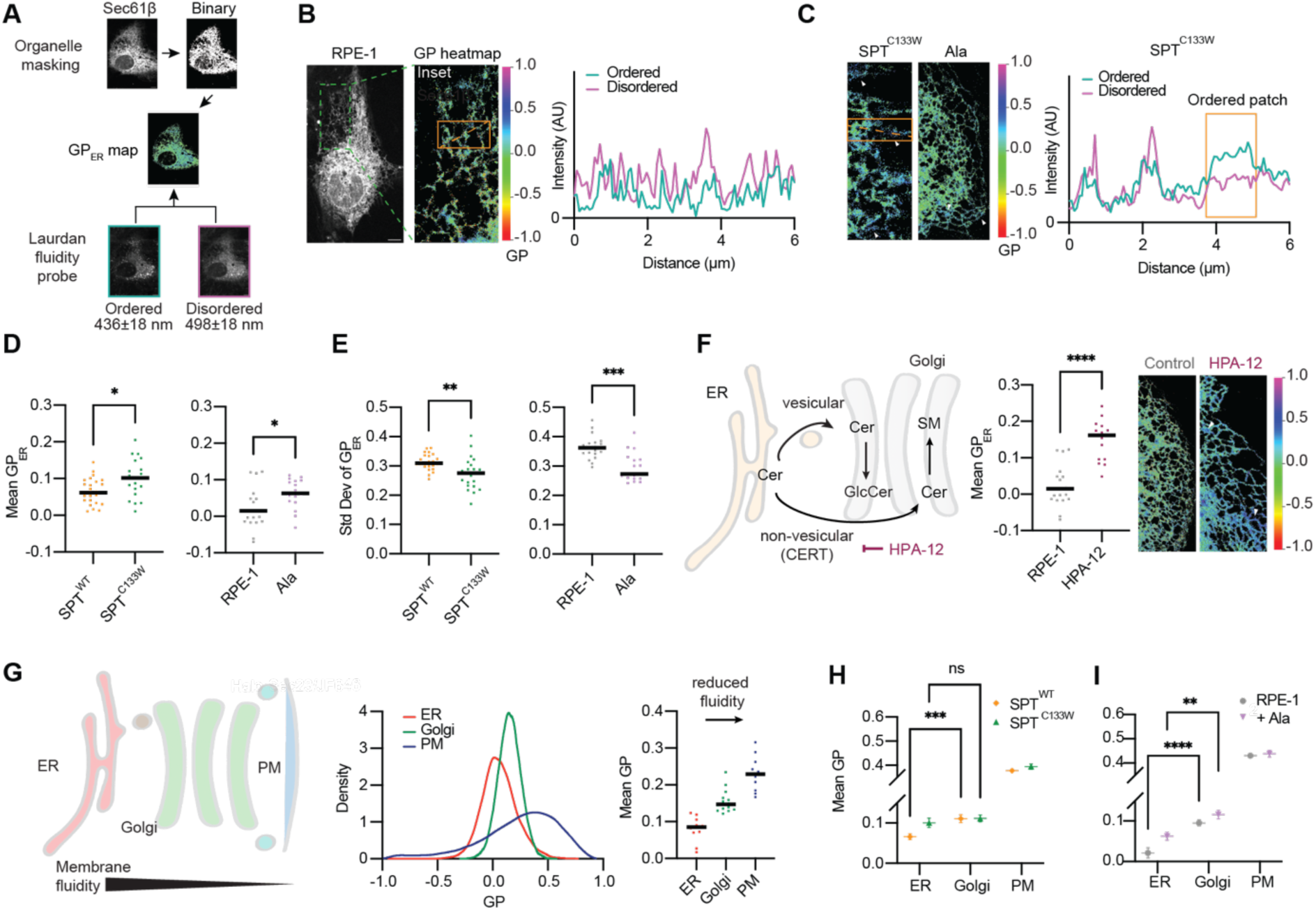
1-DeoxySL metabolism alters membrane fluidity in the secretory pathway. **A.** Methodology for generating ER-localized membrane ordering profiles using segmentation of two-channel confocal fluorescence intensity of the solvatochromic dye Laurdan. An organelle marker, Sec61β, is used to mask the ER signal. **B.** An example of GP_ER_ heatmap for an RPE-1 cell, showing distribution of the ordered and disordered channel intensities. Scale bar = 10 μm. **C.** Examples of 1-deoxySL-accumulating cells. The profile on the right shows a region of ER in an induced SPT^C133W^ cell with elevated ordered channel signal, reflecting a less fluid membrane. **D.** The mean GP_ER_ for induced SPT^C133W^ cells is higher than that for SPT^WT^ cells. Similarly, 1 mM Ala-fed RPE-1 cells show increased GP_ER_ compared to cells lacking Ala supplementation. Points represent total GP_ER_ values computed across individual cells (n = 20 (SPT^C133W^), 24 (SPT^WT^), 19 (RPE-1), 17 (Ala)). *, P < 0.05 by Mann-Whitney test. **E.** Variability in GP_ER_ within single cells is reduced in both SPT^C133W^ and Ala-fed cells, corresponding to more homogenous regions of high GP_ER_. The SD of GP_ER_ across all pixels within a single cell was computed for the same cells analyzed in D. **, P < 0.01 by Mann-Whitney test. **F.** Accumulation of canonical Cer in the ER can also reduce membrane fluidity. Cells treated with the CERT inhibitor HPA-12 show increased GP_ER_ (plot on left) and regions of high order tubules (micrographs on right). ****, P < 0.001 by Mann-Whitney test. **G.** Membrane fluidity decreases along the secretory pathway in unstressed cells, demonstrated by a monotonic increase in Laurdan GP when segmented with ER (Sec61β-mCherry), Golgi (SiT-mApple) and PM (Cell Mask Deep Red) markers. **H.** In SPT^C133W^ cells, the increase in ordering reflected between GP_ER_ and GP_Golgi_ is lost. It remains in SPT^WT^. **I.** Ala-fed cells retain an ER to Golgi GP difference, despite increases in GP_ER_. Points show mean and SEM of individual cell organelle GP in separate experiments for GP_ER_ and GP_Golgi_ and fields of cells for GP_PM_; n, SPT^WT^ = 24 (ER), 18 (Golgi), 8 (PM); n, SPT^C133W^ = 20 (ER), 15 (Golgi), 8 (PM); n, RPE-1 = 15 (ER), 12 (Golgi), 4 (PM); n, Ala = 17 (ER), 10 (Golgi), 8 (PM). **, P < 0.01; ***, P < 0.001; ****, P < 0.0001 by 2-way ANOVA.

GP-heatmaps of RPE-1 show a generally fluid ER membrane, with variable GP_ER_ values averaging 0 (**Figure 4B**). In contrast, heatmaps of induced SPT^C133W^ cells showed regions of increased ordering, which were also observed in Ala-fed cells (**Figure 4C**); both of which accumulate 1-deoxySLs. Mean GP_ER_ was increased upon SPT^C133W^ induction and Ala-feeding compared to their corresponding controls (**Figure 4D**), indicating a more ordered ER membrane. This corresponded to a reduction in variation in GP_ER_ as well, since ordered ER membrane regions showed uniformly high Laurdan GP values (**Figure 4E**). To test if the increase in GP_ER_ was due to a defect in exit of 1-deoxy products out of the ER or their intrinsic properties, we targeted distribution of canonical ceramides from the ER with HPA-12 to inhibit CERT activity^52^. Consistent with the accumulation of saturated ceramides in ER membranes, Laurdan GP_ER_ increased significantly in cells treated with HPA-12, similar to that for SPT^C133W^ cells (**Figure 4F**). Thus, retention of canonical Cers within the ER can mimic the reduction of ER membrane fluidity observed for 1-deoxySL-synthesizing cells.

Lipid composition changes along the secretory pathway due to continual enrichment of cholesterol and saturated lipids, including SLs^53^, which act to increase membrane ordering in later secretory compartments and the plasma membrane (PM). We thus extended our analysis of secretory pathway fluidity to Golgi membranes and the PM using additional markers for masking and Laurdan GP signals corresponding to these additional membranes (**Figure S6**). As expected, unperturbed RPE-1 cells showed a monotonic increase in Laurdan GP along the secretory pathway from the ER to Golgi to PM (**Figure 4G**). SPT^C133W^ cells showed an increase in GP_ER_, but did not show an increase in ordering of Golgi membranes (GP_Golgi_) or the PM (GP_PM_), consistent with the retention of 1-deoxySLs in the ER (**Figure 4H**). This caused SPT^C133W^ cells to show similar GP_ER_ and GP_Golgi_ values, while the gradient between GP_Golgi_ and GP_PM_ remained unchanged. In contrast, SPT^WT^ cells retained an ordering gradient between ER and Golgi membranes, albeit a reduced one compared to RPE-1 cells. Ala-fed cells maintained a difference between GP_ER_ and GP_Golgi_, despite an increase in the latter (**Figure 4I**). Thus, a loss of fluidity gradient in the secretory pathway might require high levels of 1-deoxySL accumulation that alterations in amino acid levels do not achieve.

In PennGreen-azide labeling, we observed that that while 1-deoxySL accumulate across the ER network, they show additional enrichment at ERES (**Figure 1D**). We thus also tested if their overproduction in SPT^C133W^ cells and subsequent accumulation at ERES could alter the morphology of these structures. To measure ERES size, we quantified the area occupied by SEC23-labeled puncta in transfected cells. We observed an increase in ERES size in SPT^C133W^ cells compared to both SPT^WT^ and RPE-1 controls (**Figure 5A**). We did not observe an increase in Ala-fed cells, suggesting another phenotype that might require high levels of ER-trapped 1-deoxySLs. We next asked if enlarged ERES results from the properties of 1-deoxySLs or their failure to traffic out of the ER. In RPE-1, we inhibited both routes for ER trafficking and processing of ceramides: the SM pathway through CERT inhibition (HPA-12), and the vesicular pathway through inhibition of glucosylceramide synthase (GCS) with 1-phenyl-2-decanoylamino-3-morpholino-1-propanol (PDMP)^54^. Lipidomics showed that HPA-12-treated cells showed only reduced accumulation of SM, while PDMP-treated cells only displayed reduced hexosyl and lactosyl ceramides, consistent with the distinct branches of post-ceramide SM metabolism (**Figure 5C**). However, only PDMP treatment caused an enlargement in ERES (**Figure 5D**), indicating that the GCS pathway in SL metabolism is tied to vesicular trafficking. This observation further supports a model in which defects in vesicular trafficking of 1-deoxySLs could drive their accumulation in ERES (**Figure 2E**) and alter these structures in cells (**Figure 5A**).

**Figure 5.**
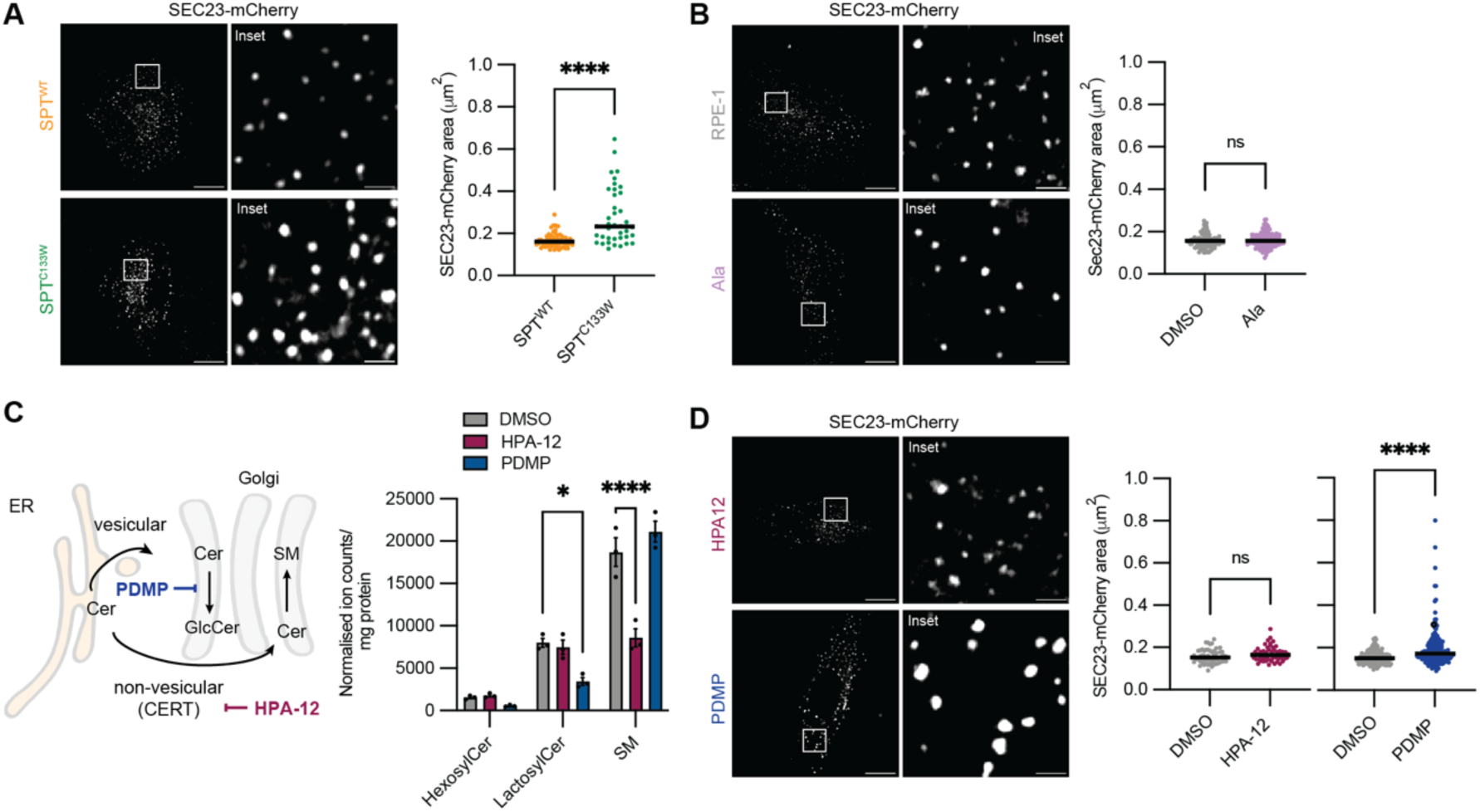
Alterations to SL metabolism alter ER exit site morphology. **A.** Individual SPT^WT^ and SPT^C133W^ cells transfected with the COPII protein SEC23-mCherry; the latter shows enlarged ERES (0.23 vs. 0.16 μm^2^) as measured by SEC23-mCherry area. For the plot on the right, each individual point represents an individual SEC23-mCherry puncta across multiple dishes; n = 60 (SPT^WT^), 40 (SPT^C133W^). **B.** Ala treatment does not enlarge the ERES of RPE-1 cells, with treated and untreated cells featuring an identical SEC23-mCherry area of 0.15 μm^2^ (RPE-1, n = 101; Ala, n = 107). **C.** Testing the roles of vesicular and non-vesicular Cer routes from the ER to Golgi. The CERT inhibitor HPA-12 inhibits non-vesicular trafficking, leading to a loss of SM levels compared to the DMSO control. The GCS inhibitor PDMP affects the vesicular trafficking route and leads to a loss of glycosylated SLs. **D.** HPA-12 treatment does not alter ERES size (mean SEC23-mCherry area of 0.16 μm^2^ in n = 59 ERES vs. 0.16 μm^2^ in n = 59 ERES of DMSO-treated control cells), while PDMP does (mean SEC23-mCherry area of 0.20 μm^2^ in n = 169 ERES vs. 0.16 μm^2^ in n = 151 ERES of DMSO-treated control cells). *, P < 0.05; ****, P < 0.001 by Mann-Whitney test. Scale bars = 11 μm (whole cell) or 1.5 μm (inset).

### 1-DeoxySLs cause cargo-specific effects on secretory trafficking

The increase in membrane ordering along the secretory pathway has been proposed to function in driving directionality and sorting of secretory cargoes at both the ER and Golgi^55–57^. Recently, it was observed that modulation of ER membrane fluidity using synthetic azobenzene-containing acyl chains caused a cargo-dependent effect on secretion rates. The release of cargoes preferring disordered membranes (TNF-α) to the Golgi was slowed when ER membrane fluidity was optically increased, while those preferring ordered membrane environments (GPI-anchored proteins) were sped up^56^. We asked if 1-deoxySLs could act through this mechanism, given the changes we observed on ER membrane fluidity. We employed the Retention Using Selective Hooks (RUSH) assay to correlate changes to secretory pathway ordering to cargo trafficking rates^58^ (**Figure 6A**). In these experiments, the addition of biotin to cells causes the dissociation of the cargoes fused to a streptavidin-binding protein (SBP) from an ER-localized ‘hook’ containing a streptavidin fused to a KDEL sequence. We focused on anterograde transport from the ER to Golgi, given that the differences across these compartments showed the largest change from our experimental perturbations.

**Figure 6.**
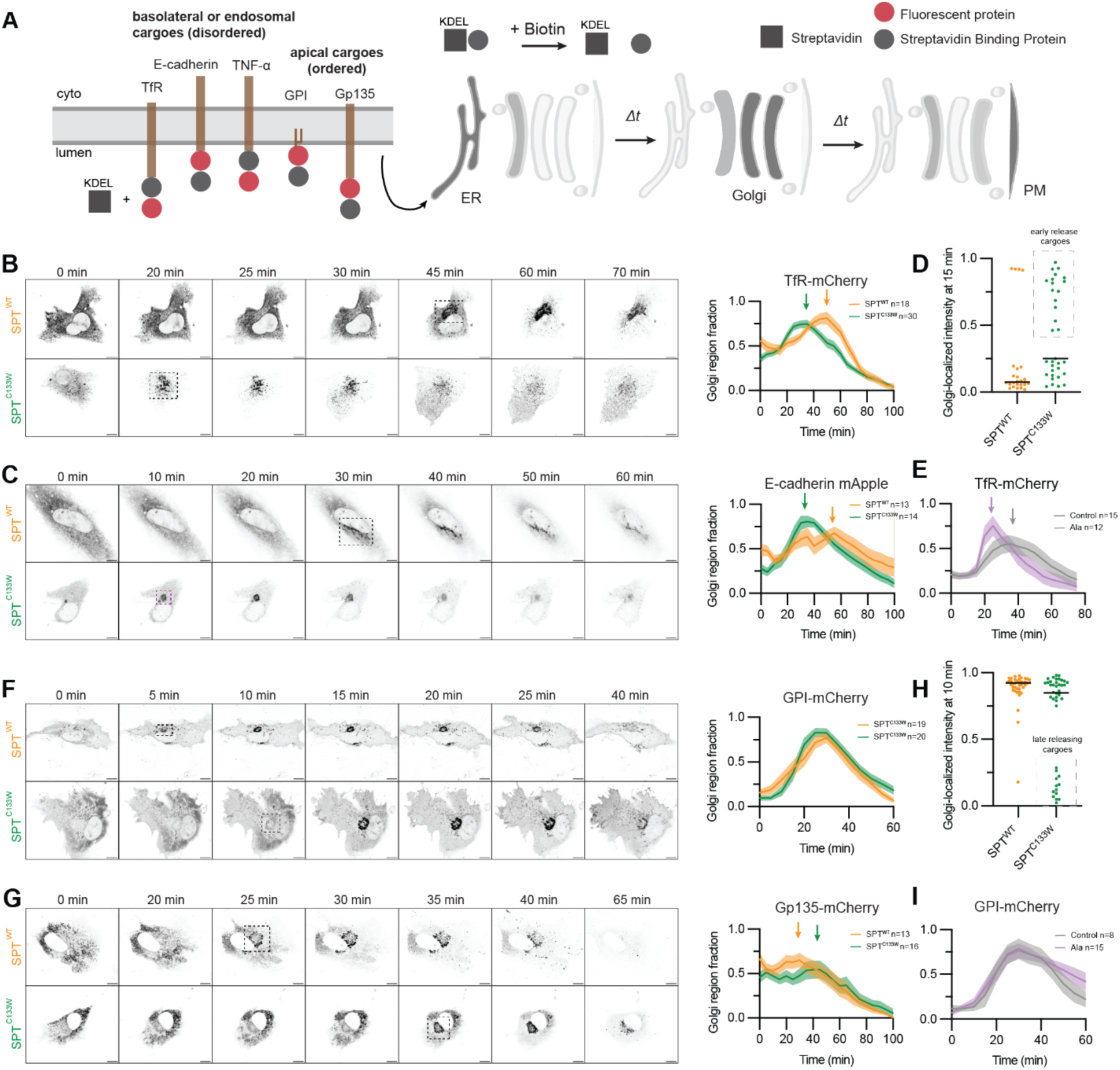
Accumulation of 1-deoxySLs modulates protein cargo release from the ER. A. Schematic of RUSH experiments for monitoring protein cargo release from the ER through the Golgi. On the left are different cargoes analyzed in this study: disordered-membrane cargoes include TfR, E-cadherin, and TNF-α, while apical membrane cargoes include GPI-anchored proteins and Gp135. **B.** Representative live cell time course comparing ER release of TfR-mCherry upon biotin addition in SPT^WT^ and SPT^C133W^ cells. The dashed box indicates the Golgi region for which fluorescence is quantified; it is shown at the time point corresponding to initial Golgi accumulation observed, which occurs earlier for SPT^C133W^. The plot on the right shows the fluorescence intensity within Golgi regions as a fraction of the whole cell. Arrows indicate times for maximum cargo concentration in the Golgi region. Number of cells (n) for each condition is provided. Scale bars = 10 μm. **C.** Similar data for the basolateral membrane protein E-cadherin-mApple. **D.** Fixed cell RUSH experiments with TfR show an early release population co-localizing with the Golgi marker in SPT^C133W^ cells 15 minutes after biotin addition. Time course data is shown in **Figure S7C**, alongside that of another disordered membrane cargo (TNF-α-mCherry). **E.** Ala-supplemented cells also show an early release of TfR-mCherry. **F.** Representative live cell time course for ER release of GPI-anchored mCherry. GPI-anchored proteins localize to apical membranes. In this case, maximum Golgi-region intensity occurs at a similar time for both SPT^WT^ and SPT^C133W^ cells, but the former show increased levels at early time points. Scale bars = 10 μm. **G.** Similar data for the apical transmembrane protein Gp135, which shows slower ER exit kinetics in SPT^C133W^ cells. **H.** Fixed cell RUSH data shows a population of SPT^C133W^ cells with unreleased GPI-mCherry at 10 minutes after biotin addition. Time course data is shown in **Figure S7D**. **I.** Ala-supplemented cells show identical GPI-mCherry ER release kinetics.

We initially analyzed the trafficking of two single-pass transmembrane proteins fused to fluorescent proteins: Transferrin Receptor (TfR-mCherry) and E-cadherin (E-cadherin-mApple) with live cell RUSH assays, in which accumulation to regions of the cell near the perinuclear-Golgi region is quantified. TfR is an internalized receptor that cycles between the PM and endosomes, while E-cadherin is cell-cell adhesion protein that localizes to basolateral membranes in polarized epithelial cells. Both these cargoes associate with liquid-disordered regions of phase-separated giant plasma membrane vesicles (GPMV), indicating an affinity for disordered membrane environments^59^ typical of intracellular compartments and basolateral plasma membranes. For both cargoes, we observed moderately more rapid trafficking from the ER to Golgi in SPT^C133W^ compared to SPT^WT^ in live cell experiments (**Figure 6B**, **Figure 6C**). We confirmed these results in fixed-cell RUSH experiments with TfR, in which Golgi-localization is assessed through co-localization with immunostained GM130 (**Figure S7**). Another disordered cargo, TNF-α-mCherry, also showed more rapid ER release in SPT^C133W^ cells in these experiments (**Figure S7C**). Since fixed cell RUSH experiments involve a larger number of cells, they allowed us to identify a fast-release population of SPT^C133W^ cells in which TfR is released within 15 minutes of biotin addition (**Figure 6D**). Ala-fed cells also showed more rapid TfR release compared to RPE-1 controls, further supporting a role for 1-deoxySLs in altering ER release kinetics for this cargo (**Figure 6E**).

We next asked whether the increased secretion rates for 1-deoxySL-producing cells was dependent on the type of cargo trafficked. We tested a pair of apical membrane cargoes, a GPI-anchored protein (GPI-mCherry)^60^ and the transmembrane protein Gp135^61^. Apical membrane proteins localize to liquid ordered regions of GPMVs^62^. The TMDs of such ‘raft-enriched’ proteins generally exit the ER faster than disordered cargoes^63^. In yeast, the sorting of GPI-anchored proteins to COPII vesicles for ER exit depends on sphingolipids^64,65^, though the extent this is true for mammalian cells is debated ^66^. For GPI-mCherry, we observed SPT^C133W^ cells did not show the enhanced secretion displayed for disordered cargoes (**Figure 6F**) and cells showed a late release population not present in SPT^WT^ (**Figure 6H**). For Gp135, SPT^C133W^ cells showed slowed release kinetics compared to SPT^WT^. Analysis of larger populations of cells at discrete timepoints in fixed cell RUSH experiments also revealed a slowed release of GPI-mCherry in SPT^C133W^ compared to SPT^WT^ and RPE-1 (**Figure S7**) and showed that sub-populations of SPT^C133W^ cells retained ER localization after release had been completed in SPT^WT^ cells (**Figure 6H**). Ala-supplemented cells, which also accumulate 1-deoxySLs, though to a lesser extent than SPT^C133W^-expressing cells, also did not show any changes to GPI-mCherry secretion rates (**Figure 6I**). In the systems tested, the accumulation of 1-deoxySLs either slows ER exit kinetics of apical membrane cargoes or leaves them unchanged. In contrast, they increased the rate of ER exit for disordered membrane cargoes.

## Discussion

1-DeoxySLs are a class of non-canonical SLs associated with, and potentially causative for, several human diseases. Since their discovery as disease-associated lipid class, there have been efforts to identify cytotoxic roles for 1-deoxySLs. While such roles are certainly possible, the tissue-specific nature of genetic disorders caused by Ala-preferring SPT mutations suggest that more nuanced effects could also be relevant for their pathology. Answering these questions is challenging because the fundamental mechanisms by which 1-deoxySLs are metabolized and transported are unknown, as are their effects on broader cellular functions. Here we used a combination of approaches to better understand the role of 1-deoxySLs in an RPE cell line that is a commonly used model for secretory pathway biology. We found that 1-deoxy products are largely retained in the ER, where they alter the structure and membrane properties of this organelle. Although these effects do not elicit ER stress response, they can affect rates at which protein cargoes are released to Golgi compartments in the secretory pathway. Some of the effects of 1-deoxySL metabolism (reduced ER membrane fluidity, increased release of Transferrin receptor) were observed both from increases in Ala concentration and from expression of a disease-associated promiscuous SPT variant, while more drastic effects such as enlarged ERES and inhibited release of apical membrane cargoes only occurred during the latter. These discrepancies could result from the larger increase in 1-deoxySLs due to reduced L-serine selectivity in SPT^C133W^ than for changes in amino acid concentrations (**Figure 3**), although we cannot rule out other mechanisms by which they might act. As a method for investigating 1-deoxySL biology, expression of mutant SPT alleles overcomes the pleiotropic effects of bulk amino acid changes while avoiding the toxicity and extreme overproduction caused by supplementation with high amounts of 1-deoxySA.

In the canonical sphingolipid pathway, Cer is transported from the ER to the Golgi, where it is processed in complex SLs. This occurs via at least two known pathways. The first is through vesicular trafficking at ERES, the site of protein secretion, to the *cis*-Golgi. Vesicular secretion of ceramides is better characterized in yeast, where it is predominant, than mammalian cells^43^. The second is through non-vesicular transport via the LTP CERT, which acts at contact sites between the ER and the *trans*-Golgi and predominantly supplies Cer for SM synthesis^39^. We initially suspected that CERT-mediated transport of 1-deoxyCer could be impeded, since the START domain of CERT makes direct polar contacts with the C1-hydroxyl of Cer. However, we found that recombinant CERT START domain is able to transport a model 1-deoxySL *in vitro*, and in fact does so at a higher rate than its corresponding DHCer species. Two additional lines of evidence support that vesicular trafficking of 1-deoxySL could be impaired. First, organelle-specific labeling of an alkyne 1-deoxySL metabolic probe shows extensive accumulation at or near ERES, indicating that they might not be efficiently trafficked through anterograde carriers. Second, overproduction of 1-deoxySLs by SPT^C133W^ cells causes an enlargement of ERES visualized by the COPII protein SEC23. We observed similar effects in non-1-deoxySL producing cells whose glucosylceramide synthase is inhibited, reflecting a potential backup of canonical Cer trafficking through ERES to the *cis*-Golgi. Limited vesicular trafficking of 1-deoxySLs could result from effects on anterograde vesicle formation itself or because of inability of glycoSL metabolism to subsequently modify 1-deoxy products. In contrast, cells with inhibited CERT did not show enlarged ERES, consistent with the location of non-vesicular trafficking of Cers occuring at ER *trans*-Golgi contact sites, not at ERES. However, they did show increased ER membrane ordering, reflecting a build-up of saturated Cer in the ER.

Even though CERT transports 1-deoxyDHCer *in vitro*, non-vesicular transport of 1-deoxy cargoes might still be limited in cells. We observed that 1-deoxyCer and 1-deoxyDHCers produced in RPE-1 primarily contain VLCFA *N*-acyl chains greater than 20 carbons in length. Enrichment of VLCA chains in 1-deoxy species had been previously reported in RAW macrophages^7^ and HCT116 colorectal cancer cells^31^, suggesting that it is a general feature determined by the differing substrate specificities of ceramide synthases (CerS), like CerS2^67^ and CerS5^68^. VLCFA-containing Cers are less efficiently transported by CERT^69^, which could underlie the role of CERT in mediating their transport to the *trans*-Golgi for conversion into short N-acyl chain SM by SMS1. In contrast, it was proposed that VLCFA-containing ceramides are sorted into vesicular carriers for ER export^70^, where they are delivered to the *cis*-Golgi for GCS. Our data suggest that 1-deoxyCer and 1-deoxyDHCer could operate under similar constraints. Thus, the metabolism of both SLs and 1-deoxySLs might directly impact their trafficking patterns in cells.

The lack of a polar group on the sphingoid backbone of 1-deoxySLs alters their biophysical properties, namely their hydrophobicity and immiscibility with other lipids. Dihydro species, which do not accumulate in canonic SLs but do in 1-deoxys, show even more extreme properties. Indeed, in our experience, both 1-deoxyCers and 1-deoxyDHCers are challenging to reconstitute into synthetic liposomes using standard protocols. Additionally, cell-synthesized 1-deoxySLs are enriched in VLCFA *N*-acyl chains, which would further contribute to these properties. The ER membrane is predominantly composed of unsaturated phospholipids, leading to a highly fluid membrane that can support rapid protein and lipid dynamics. In this context, the accumulation of long-chain saturated or monounsaturated Cer/DHCer species could dramatically alter membrane properties. Consistent with this hypothesis, we observe reduced levels of membrane fluidity in 1-deoxySL producing cells. In our analysis, it is unclear if this reflects simply a more ordered membrane environment, or the formation of localized ordered domains. The Laurdan dye we use shows low fluorescence in solid, gel-like domains^71^, which ceramides^72^ and their 1-deoxy analogues^8^ readily form, so it is possible that our analysis does not fully capture these effects. The increase in ER Laurdan GP (reduced fluidity) caused by 1-deoxySL synthesis can be replicated by inhibition of canonical Cer trafficking (via CERT inhibition). Thus, the retention of 1-deoxySL could drive biophysical changes to ER membranes independently of the unique properties of these lipids that are not present in canonical Cers.

In cells that accumulate large amounts of 1-deoxyCer and 1-deoxyDHCer, ER membranes show similar membrane ordering, measured with Laurdan GP, as Golgi membranes that follow in the secretory pathway. A gradient of membrane fluidity and thickness across the secretory pathway has long been proposed to function in membrane protein trafficking and sorting, including to the Golgi^73^. A wide range of studies have shown that membrane domains or anchors of secreted proteins could be matched to their host lipid bilayer to maintain proper sorting during secretion^63,74–76^. In polarized cells, PM proteins are thought to segregate by a similar mechanism, potentially at endosomes^77^, with basolateral proteins matching with more disordered membranes and apical proteins with more ordered, raft-like membranes^78,79^. The latter are enriched in sphingolipids and cholesterol^80^. While comparative studies on different membrane proteins have shed light on their secretory trafficking, less has been done to ask how lipid perturbations affect this process due to the challenges in manipulating lipid composition. Much of the focus in this area has been on post-Golgi trafficking and sorting of apical and basolateral membrane proteins^81,82^. More recently, disruption of the ER-*trans*-Golgi cholesterol transporter carried out by oxysterol binding protein (OSBP) alters apical vs. basolateral cargo secretion and sorting at the Golgi in polarized cells^83^. Much like SLs, cholesterol forms an increasing gradient along the secretory pathway, despite its synthesis in the ER due to highly active export pathways^84^.

Cargo export from the ER itself could also be dependent on changes in secretory pathway membrane ordering, as suggested by reported effects on reduced SL synthesis inhibition upon myriocin treatment^56^ or knockdown of ether lipid synthesis^85^. One model for these effects is that cargo sorting into the ERES itself is dependent on its bilayer mismatch with the rest of the ER matrix. Apical cargoes, like GPI-anchored proteins, show a bilayer mismatch with the disordered ER membrane, which could be relevant for their rapid sorting into ERES enriched in sterols and sphingolipids better reflecting the plasma membrane viscosity. Through this mechanism, an increase in ER membrane ordering caused by failure of 1-deoxyCer and 1-deoxyDHCer to exit the ER could reduce cargoes affinity for ERES foci, leading to higher diffusion through the ER network. In contrast, cargoes tailored for thinner and more disordered membranes may not match with more ordered ER membranes caused by 1-deoxySLs, accelerating their sorting toward ERES. Such a model depends on the existence of multiple classes of exit carriers for different types of cargoes, which has long been hypothesized^86,87^. Although measuring the membrane microenvironment of ERES themselves remains challenging, recent work showing they are interconnected with the *cis*-Golgi suggest that they might approximate the more ordered state of that compartment^44^. Notably, the difference between membrane ordering of the ER and Golgi (GP_ER_ and GP_Golgi_) is lost in 1-deoxySL-synthesizing cells.

Given their substantial effects on the packing of lipid membranes and their disparate membrane concentrations, we propose that the dysregulation of SL homeostasis could be especially relevant for the processing of proteins through the secretory pathway. The pathophysiology of SL-related diseases, including those that affect degradation^88^, and trafficking^89^, remains mysterious. In the case of diseases that are thought to be driven by increased 1-deoxy bases, effects on trafficking of specific membrane protein cargoes could be relevant for understanding their cellular targets. For MacTel, sorting of proteins is required to support the polarized proteome of RPE that support photoreceptors. In HSAN1, axonal neuropathy is accompanied by the loss of myelin, a structure enriched in secreted SLs and membrane proteins with affinities for ordered membrane^90^. Retention of 1-deoxySL products within early secretory compartments could disrupt both these processes and contribute to cell type-specific pathologies that depend on trafficking of individual membrane proteins synthesized in the ER. Similar dynamics could be relevant for other diseases related to the SL metabolism and transport.

**Schematic:**
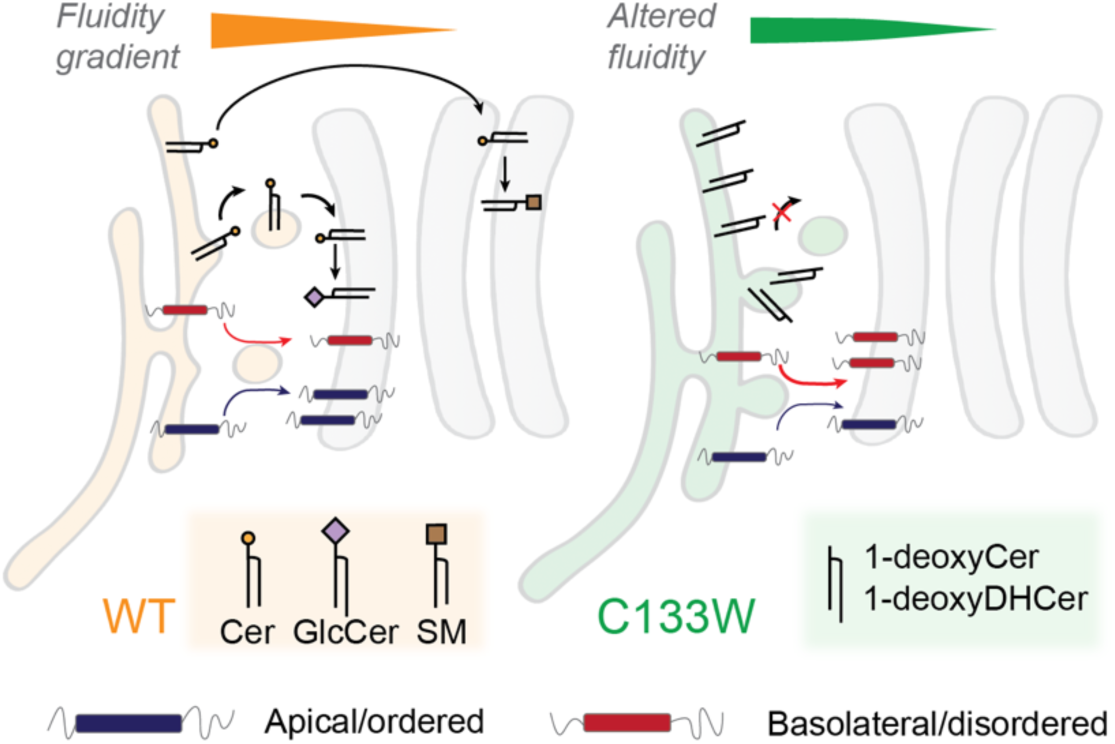
Model for dysregulation of early secretory trafficking by 1-deoxySL metabolism. With WT SPTLC-1, ceramides are trafficked from the ER to the *trans*-Golgi for SM synthesis via CERT, or via vesicles to the *cis*-Golgi for GlcCer synthesis. This lipid trafficking alongside that of cholesterol drives a monotonic membrane fluidity or ordering gradient along secretory compartments. Cargo trafficking of disordered and ordered membrane proteins is balanced. For cells expressing mutant SPTLC-1 variants like C133W, 1-deoxySLs accumulate due to inefficient trafficking out of the ER. This increases ER membrane ordering and alters cargo release from the ER to Golgi.

## Acknowledgments

Dorota Skowronska-Krawczyk, Chris Obara, Aubrey Weigel, Ivan Castello-Serrano, and David Kovács provided helpful discussions. Kailash Venkatraman provided experimental support. Dorota Skowronska-Krawczyk and Jennifer Lipincott-Schwartz provided reagents. Funding was provided by the National Institutes of Health (R35-GM142960 to I.B., R01-CA234245 to C.M.M., R35-GM141939 to N.K.D., R35-GM133565 to S.N., T32-GM146648 to A.W.), the National Science Foundation (MCB-2047391 to S.N.), HHMI (Freeman Hrabowski Scholar Program to S.N.), MCIN/AEI/10.13039/501100011033 (R.J.B.), and the European Regional Development Fund (RYC2020-030065-I to R.J.B.).

## Author contributions

Y-T.T., C.M.M., and I.B. conceived the project. Y-T.T., N-F.L., O.S., A. L., K.S-K., and G.H.M carried out experiments. R.V. analyzed data. A.W., O.S., and R.J.B. synthesized reagents. N.K.D., L.K., S.N., C.M.M., and I.B. supervised the project and acquired funding.

## Declaration of interests

The authors declare no competing interests.

## Materials and methods

### Synthesis of PennGreen-azide

Commercially available *N*-phenyl-bis (trifluoromethanesulfonimide), 1,4-dioxane, triethylamine (Et_3_N), lithium iodide (LiI), *N*-Boc-ethylenediamine, palladium(II) acetate [Pd(OAc)_2_], cesium carbonate (Cs_2_CO_3_), 4,5-Bis(diphenylphosphino)-9,9-dimethylxanthene (Xantphos), trifluoroacetic acid (TFA), O-(7-azabenzotriazol-1-yl)-1,1,3,3-tetramethyluronium hexafluorophosphate (HATU), *N*,*N*-diisopropylethylamine (DIEA), and formic acid were obtained from Sigma-Aldrich. 2,7-Difluoro-6-hydroxy-9-(2-methylphenyl)-*3H*-xanthen-3-one (Pennsylvania Green) was obtained from AK Scientific, Inc. Azidoacetic acid was obtained from TCI Chemicals. Deuterated chloroform (CDCl_3_) was obtained from Cambridge Isotope Laboratories. All reagents obtained from commercial suppliers were used without further purification unless otherwise noted. Analytical thin-layer chromatography was performed on E. Merck silica gel 60 F_254_ plates. Silica gel flash chromatography was performed using E. Merck silica gel (type 60SDS, 230-400 mesh). Solvent mixtures for chromatography are reported as v/v ratios. HPLC analysis was carried out on an Eclipse Plus C8 analytical column with *Phase A*/*Phase B* gradients [*Phase A*: H_2_O with 0.1% formic acid; *Phase B*: MeOH with 0.1% formic acid]. HPLC purification was carried out on Zorbax SB-C18 semipreparative column with *Phase A*/*Phase B* gradients [*Phase A*: H_2_O with 0.1% formic acid; *Phase B*: MeOH with 0.1% formic acid]. Proton nuclear magnetic resonance (^1^H NMR) spectra were recorded on a Varian VX-500 MHz MHz spectrometer and were referenced relative to residual proton resonances in CDCl_3_ (at δ 7.24 ppm). ^1^H NMR splitting patterns are assigned as singlet (s), doublet (d), triplet (t), quartet (q) or pentuplet (p). All first order splitting patterns were designated on the basis of the appearance of the multiplet. Splitting patterns that could not be readily interpreted are designated as multiplet (m) or broad (br). Carbon nuclear magnetic resonance (^13^C NMR) spectra were recorded on a Varian VX-500 MHz spectrometer and were referenced relative to residual proton resonances in CDCl_3_ (at δ 77.23 ppm). Electrospray Ionization-Time of Flight (ESI-TOF) spectra were obtained on an Agilent 6230 Accurate-Mass TOFMS mass spectrometer.

#### 2,7-difluoro-6-iodo-9-(*o*-tolyl)-*3H*-xanthen-3-one (PennGreen iodide)^1^

To a dry, argon-flushed round-bottomed flask was added Pennsylvania Green^2,3^ (PennGreen, 50.0 mg, 147.8 µmol) and *N*-phenyl-bis(trifluoromethanesulfonimide (63.4 mg, 177.4 µmol). Then, anhydrous 1,4-dioxane (1 mL) was added, followed by Et_3_N (24.7 µL, 177.4 µmol). The flask was heated under Ar to 60 °C for 1 h [Note: Conversion to the intermediate triflate was observed by TLC and HPLC-MS]. The flask was removed from the oil bath and LiI (59.3 mg, 443.4 µmol) and a reflux condenser were added. The solution was refluxed (∼110 °C) for 4 h. The flask was cooled to rt. Then, aqueous 2M solution of NaOH (250 µL) was added, and the solution was stirred for 1 h. Afterwards, H_2_O (3 mL) was added, giving rise to a product slurry that was stirred for 1 h. After this time, the orange slurry was chilled to 4 °C, filtered, and the product cake washed with cold H_2_O (3 × 1 mL) to give the product containing residual 1,4-dioxane. This product cake was treated with Et_2_O (250 µL), stirred vigorously for 1 min, and hexanes (250 µL) were added. The resulting slurry was filtered and the product cake extensively dried under high vacuum to provide 53.8 mg of PennGreen iodide as an orange solid [81%].^1^H NMR (CDCl_3_, 500.13 MHz, δ): 7.97 (d, *J_1_* = 5.1 Hz, *J_2_* = 0.8 Hz, 1H, 1 × CH), 7.52 (td, *J_1_* = 7.6 Hz, *J_2_* = 1.4 Hz, 1H, 1 × CH), 7.42 (ddd, *J_1_* = 16.4 Hz, *J_2_* = 7.9 Hz, *J_3_* = 1.4 Hz, 2H, 2 × CH), 7.14 (dd, *J_1_* = 7.6 Hz, *J_2_* = 1.4 Hz, 1H, 1 × CH), 6.70 (d, *J_2_* = 7.8 Hz, 1H, 1 × CH), 6.62 (dd, *J_1_* = 10.5 Hz, *J_2_* = 0.9 Hz, 1H, 1 × CH), 6.56 (dd, *J_1_* = 6.8 Hz, *J_2_* = 1.3 Hz, 1H, 1 × CH), 2.07 (s, 3H, 1 × CH_3_). ^13^C NMR (CDCl_3_, 125.77 MHz, δ): 176.9 (d, *J* = 21.2 Hz), 159.4, 157.9, 157.4, 156.9, 155.8, 153.8, 148.1, 136.1, 131.4, 131.2, 130.4, 129.0, 128.0 (d, *J* = 2.1 Hz), 126.8, 121.4 (dd, *J*_1_ = 48.8 Hz, *J*_2_ = 8.1 Hz), 111.1 (dd, *J*_1_ = 267.8 Hz, *J*_2_ = 25.0 Hz), 107.1 (d, *J* = 4.6 Hz), 87.9 (d, *J* = 29.2 Hz), 67.2,* 19.8. MS (ESI-TOF) [m/z (%)]: 448 ([MH]^+^, 100).* NMR artifact (1,4-dioxane).

#### 2,7-difluoro-6-(*N*-Boc-ethylenediamine)-9-(*o*-tolyl)-*3H*-xanthen-3-one (*N*-Boc-EthD-PennGreen)

An oven-dried microwave test tube was charged with PennGreen iodide (10.0 mg, 22.3 µmol), *N*-Boc-ethylenediamine (5.3 µL, 33.5 µmol), Pd(OAc)_2_ (0.5 mg, 2.2 µmol), Cs_2_CO_3_ (14.5 mg, 44.6 µmol), and Xantphos (1.9 mg, 3.35 µmol). The test tube was slowly flushed with Ar. Then, anhydrous, degassed toluene (500 µL) was added. The test tube was sealed and heated to 120 °C in a microwave reactor for 1 h. The reaction tube was cooled to rt and diluted with CH_2_Cl_2_ (2.5 mL). Then, H_2_O (1.5 mL) was added, and the organic layer was separated. The aqueous portion was extracted with CH_2_Cl_2_ (2 × 1.5 mL). The organic layers were combined, dried over anhydrous Na_2_SO_4_, and filtered. The solvent was removed under reduced pressure, giving an orange crude solid. The crude was purified by flash chromatography (0-5% MeOH in CH_2_Cl_2_), affording 6.1 mg of *N*-Boc-EthD-PennGreen as a red solid [57%, R_f_ = 0.16 (2% MeOH in CH_2_Cl_2_)]. ^1^H NMR (CDCl_3_, 500.13 MHz, δ): 7.48 (td, *J_1_* = 7.5 Hz, *J_2_* = 1.4 Hz, 1H, 1 × CH), 7.44-7.34 (m, 2H, 2 × CH), 7.13 (dd, *J_1_* = 7.6 Hz, *J_2_* = 1.4 Hz, 1H, 1 × CH), 6.80-6.53 (m, 4H, 4 × CH), 5.03-4.88 (m, 1H, 1 × NH), 4.11-3.99 (m, 1H, 1 × NH), 3.60-3.50 (m, 2H, 1 × CH_2_), 3.49-3.39 (m, 2H, 1 × CH_2_), 2.06 (s, 3H, 1 × CH_3_), 1.45 (s, 9H, 3 × CH_3_). ^13^C NMR (CDCl_3_, 125.77 MHz, δ): 174.6, 157.6, 157.1, 154.8, 152.9, 152.0, 149.9, 148.0, 144.1, 136.0, 132.3, 131.0, 130.1, 128.9, 126.5, 115.2, 111.4 and 111.2, 110.6, 106.3, 96.8, 80.7, 39.5, 29.8, 28.4, 19.8. MS (ESI-TOF) [m/z (%)]: 481 ([MH]^+^, 100).

#### 2,7-difluoro-6-(ethylenediamine)-9-(*o*-tolyl)-*3H*-xanthen-3-one (H_2_N-EthD-PennGreen)

A solution of *N*-Boc-EthD-PennGreen (3.0 mg, 6.3 μmol) in 200 µL of TFA/CH_2_Cl_2_ (1:1) was stirred at rt for 15 min. After removal of the solvent, the residue was dried under high vacuum for 3 h and used without further purification. MS (ESI-TOF) [m/z (%)]: 381 ([MH]^+^, 100).

#### 2,7-difluoro-6-(azidoacetylethylenediamine)-9-(*o*-tolyl)-*3H*-xanthen-3-one (N_3_-Ac-EthD-PennGreen; PennGreen-azide)

A solution of azidoacetic acid (0.5 µL, 6.3 μmol) in 100 µL of CH_2_Cl_2_/DMF (1:1) was stirred at 0 °C for 10 min, and then HATU (2.6 mg, 6.9 μmol) and DIEA (4.4 µL, 25.0 µmol) were successively added. After 10 min stirring at 0 °C, a H_2_N-EthD-PennGreen (2.4 mg, 6.3 μmol) solution in 100 µL of CH_2_Cl_2_/DMF (1:1) containing DIEA (2.2 µL, 12.5 µmol) was added. After 1 h stirring at rt, the solvent was removed under reduced pressure to give a red orange solid. The corresponding residue was dissolved in MeOH (200 µL), filtered using a 0.2 µm syringe-driven filter, and the crude solution was purified by HPLC, affording 2.4 mg of N_3_-Ac-EthD-PennGreen (PennGreen Azide) as an orange solid [84%, t_R_ = 8.1 min (Zorbax SB-C18 semipreparative column, 50% *Phase A* in *Phase B*, 1 min, then 50-5% *Phase A* in *Phase B*, 5 min, and then 5% *Phase A* in *Phase B*, 7 min)]. ^1^H NMR (CDCl_3_, 500.13 MHz, δ): 7.51 (td, *J_1_* = 7.7 Hz, *J_2_* = 1.4 Hz, 1H, 1 × CH), 7.41 (dd, *J_1_* = 16.0 Hz, *J_2_* = 7.8 Hz, 2H, 2 × CH), 7.35-7.28 (m, 1H, 1 × CH), 7.16-7.10 (m, 1H, 1 × CH), 7.07 (br s, 1H, 1 × NH), 6.92 (d, *J* = 6.8 Hz, 1H, 1 × CH), 6.73 (dd, *J_1_* = 11.0 Hz, *J_2_* = 5.0 Hz, 2H, 2 × CH), 6.57 (br s, 1H, 1 × NH), 4.06 (s, 2H, 1 × CH_2_), 3.84-3.65 (m, 2H, 1 × CH_2_), 3.64-3.55 (m, 2H, 1 × CH_2_), 2.04 (s, 3H, 1 × CH_3_). ^13^C NMR (CDCl_3_, 125.77 MHz, δ): 169.3, 166.0, 161.5, 156.6, 153.8, 150.3, 148.4, 135.9, 131.9, 131.1, 130.3, 130.2, 128.8, 128.0, 126.5, 115.1, 111.6 and 111.4, 111.2, 106.1, 97.0, 52.5, 44.9, 38.6, 19.8. MS (ESI-TOF) [m/z (%)]: 464 ([MH]^+^, 100).

### Synthesis of C12-NBD-DHCer and C12-NBD-deoxyDHCer

Commercially available 2S-amino-1,3R-octadecanediol and 2S-amino-3R-octadecanol were obtained from Cayman Chemical. NBD-dodecanoic acid was obtained from Avantor Sciences. Hydroxybenzotriazole (HOBt), O-(7-azabenzotriazol-1-yl)-1,1,3,3-tetramethyluronium hexafluorophosphate (HATU), N,N-diisopropylethylamine (DIEA), and formic acid were obtained from Sigma-Aldrich. Solvent mixtures for chromatography are reported as volume/volume (v/v) ratios. HPLC analysis was carried out on a 1260 Infinity II LC System equipped with an Agilent C18 column, using Phase A/Phase B gradients (Phase A: H₂O with 0.1% trifluoroacetic acid; Phase B: acetonitrile (ACN) with 0.1% trifluoroacetic acid). Proton nuclear magnetic resonance (^1^H NMR) spectra were recorded on a 400 MHz Varian Mercury Plus spectrometer and referenced relative to residual proton resonances in CDCl₃ (δ 7.26 ppm). Carbon nuclear magnetic resonance (^13^C NMR) spectra were recorded on the same instrument and referenced to CDCl₃ (δ 77.36 ppm). High-resolution mass spectrometry (HR-MS) measurements were obtained using an Agilent 1260 Infinity Binary LC instrument coupled to a 6230 Accurate-Mass TOFMS system.

#### N-((2R)-1,3-dihydroxyoctadecan-2-yl)-12-((7-nitrobenzo[c][1,2,5]oxadiazol-4-yl)amino)dodecanamide (C12-NBD-DHCer)

NBD-dodecanoic acid (3.8 mg, 10.1 µmol) was dissolved in 200 µL of DMF in a glass vial. HATU (4.8 mg, 12.6 µmol), DIEA (13.9 µL, 80.7 µmol), and HOBt (1.4 mg, 10.6 µmol) were added successively. The mixture was stirred under N₂ for 5 minutes at room temperature (RT), then a solution of 2S-amino-1,3R-octadecanediol (3.1 mg, 10.2 µmol) in 200 µL of DMF was added dropwise. The reaction mixture was stirred overnight at RT under N₂. After solvent removal in vacuo, an orange oil was obtained, which was dissolved in methanol, filtered through a 0.2 µm syringe-driven filter, and purified by HPLC. The product was isolated as a yellow oil (5.0 mg, 78% yield; t_R_= 8.5 min; Agilent C18 semipreparative column; 50:50 Phase A:B for 2 min, ramp to 25:75 over 8 min, then to 1:99 over 4 min). ¹H NMR (CDCl₃, 400 MHz, δ): 8.54 (d, J = 9.0 Hz, 1H, 1 × CH), 6.47 (d, J = 9.0 Hz, 1H, 1 × CH), 4.15 (m, 1H, 1 × CH), 4.02 (m, 1H, 1 × CH), 3.79 (m, 2H, 1 × CH₂), 3.49 (t, J = 4.0 Hz, 2H, 1 × CH₂), 2.26 (t, J = 7.6 Hz, 2H, 1 × CH₂), 1.81–1.64 (m, 4H, 2 × CH₂), 1.34–1.21 (br m, 32H, 16 × CH₂), 0.88 (t, J = 7.0 Hz, 3H, 1 × CH₃). ¹³C NMR (CDCl₃, 101 MHz, δ): 144.1, 134.2, 124.2, 77.1 (solvent), 61.0, 39.0, 36.0, 33.7, 32.0, 30.0, 29.8, 29.6, 29.5, 29.4, 27.2, 26.1, 22.8, 14.2. HR-ESI-TOFMS: *m/z* [M + Na]^+^ calcd for C_36_H_63_N₅O_6_Na = 684.4671; found = 684.4670.

#### N-((2R)-3-hydroxyoctadecan-2-yl)-12-((7-nitrobenzo[c][1,2,5]oxadiazol-4-yl)amino)dodecanamide (C12-NBD-deoxyDHCer)

NBD-dodecanoic acid (3.8 mg, 10.1 µmol) was dissolved in 200 µL of DMF in a glass vial. HATU (4.8 mg, 12.6 µmol), DIEA (13.9 µL, 80.7 µmol), and HOBt (1.4 mg, 10.6 µmol) were added successively. The mixture was stirred under N₂ for 5 minutes at RT, then a solution of 2S-amino-3R-octadecanol (2.9 mg, 10.2 µmol) in 200 µL of DMF was added dropwise. The reaction mixture was stirred overnight at RT under N₂. After solvent removal in vacuo, a yellow oil was obtained, which was dissolved in methanol, filtered through a 0.2 µm syringe-driven filter, and purified by HPLC. The product was isolated as a orange oil (6.1 mg, 91% yield; t_R_= 8.6 min; Agilent C18 semipreparative column; 50:50 Phase A:B for 2 min, ramp to 25:75 over 8 min, then to 1:99 over 4 min). ¹H NMR (CDCl₃, 400 MHz, δ): 8.54 (d, 1H, 1 × CH), 6.19 (m, 1H, 1 × CH), 4.02 (m, 2H, 1 × CH₂), 3.65 (m, J = 6.4 Hz, 1H, 1 × CH), 3.49 (d, J = 6.4 Hz, 2H, 1 × CH₂), 2.21 (t, J = 7.6 Hz, 2H, 1 × CH₂), 1.89–1.60 (m, 6H, 3 × CH₂), 1.45–1.20 (br m, 28H, 14 × CH₂), 0.87 (t, J = 6.8 Hz, 3H, 1 × CH₃). ¹³C NMR (CDCl₃, 101 MHz, δ): 144.1, 136.4, 134.6, 73.97, 50.7, 33.3, 31.8, 29.5, 29.5, 29.5, 29.4, 29.4, 29.4, 29.3, 29.2, 29.1, 29.1, 29.0, 29.0, 28.9, 28.9, 28.3, 26.7, 25.8, 25.6, 22.5, 14.0. HR-ESI-TOFMS: *m/z* [M + Na]^+^ calcd for C_36_H_63_N₅O_5_Na = 668.4721; found = 668.4727.

### Cell culture

hTERT-RPE-1 cells (ATCC CRL-4000) were cultured in DMEM/F12 medium (Gibco) with L-glutamine and 15mM HEPES, supplemented with 10% dialyzed fetal bovine serum (Gibco) and 1% penicillin-streptomycin (Gibco). Cells were maintained in a 37°C humidified incubator kept in 5% CO_2_ atmosphere. Cells were routinely tested for mycoplasma by Human Embryonic Stem Cell Core (Sanford Consortium). For microscopy, cells were seeded into 35 mm glass bottom dishes #1.5 coverslip (MatTek) coated with 2 μg fibronectin/cm^2^ (Sigma-Aldrich). For transient expression of proteins, cells were transfected with Lipofectamine 3000 (Invitrogen) in Opti-MEM media (Gibco) for 5 h and changed into complete growth medium. Cells were imaged 24 h after transfection or later depending on the treatment.

### Confocal microscopy

All cells were grown and transfected on MatTek dishes with #1.5 coverslip. Confocal microscopy was performed on a Zeiss LSM 880 microscope equipped with plan-apochromat 64x/1.4 NA or 20x/0.8 NA objectives. For cells expressing Halo-tagged proteins and incubated with JF646 ligand, a 633 nm HeNe laser was used at 1-2% power. For Pennsylvania Green experiments, cells were excited with a 488 nm Argon laser at 0.5% power. For Cy3, mCherry, and mApple experiments, a 561 nm diode laser was used. All images used for intensity quantification were acquired in LSM mode. Airyscan imaging was performed using Plan Apo 63x/1.4 objective on ZEISS LSM 880 with Airyscan. Live-cell confocal experiments were conducted with cells incubated at 37°C with 5% CO_2_ and humidified air. For GP measurements, Laurdan dye was excited with 405 nm diode laser set at 0.3% power and emission was detected using QUASAR GaAsP detector set to two simultaneous spectrum windows: 436±18 for ordered membrane emission and 498±18 for disordered membrane emission.

### Image analysis

All image analysis was performed using Python and ImageJ (National Institutes of Health). The following Python packages were used in computation: numpy, pandas, skimage, seaborn, matplotlib and os. Specific thresholds were applied to binarize the image in Python, essentially creating a mask for the fluorescent region of interest.

### Statistical analysis

Data were statistically analyzed using GraphPad Prism 10.1.1 (GraphPad Software). Specific statistical tests were chosen according to the experiment design and specified on the text.

### Bioorthogonal labeling

The alkyne-sphinganine (alkyne-SA) and alkyne-deoxysphinganine (alkyne-deoxySA) probes were previously described^24^. Cy3-azide was purchased from Sigma-Aldrich (Cat# 777315). For Cu(I)-catalysed azide-alkyne cycloadditions (CuAAC), cells were fed with 0.1 μM of either alkyne-SA or alkyne-deoxySA for 17 hr, after which cells were rinsed with PBS and fixed with cold 4% paraformaldehyde and 0.1% glutaraldehyde in PBS for 20 min. Fixed cells were quenched with 100 mM glycine in PBS, washed twice, and CuAAC was performed by adding a mixture of 5 mM PennGreen-azide or Cy3-azide, 100 mM CuSO_4_, 500 mM BTTAA and 1 mM ascorbic acid in PBS. Cells were allowed to react for 45 min in the dark at room temperature, followed by 3 washes with PBS and stained with Hoescht (1:5,000 dilution) for 5 min. Additional 3 washes were carried out before imaging. For colocalization experiments with ERES (Figure 4), cells were transfected with SEC23-mCherry. To verify the localization of PennGreen-azide probe (Figure S2A), cells were transfected with Sec61β-mCherry (ER) or SiT-mApple (Golgi) and also stained with either MitoTracker Red FM (Thermo Fisher Scientific) or LysoTracker Deep red (Thermo Fisher Scientific). To quantify the fluorescent intensity of each PennGreen or Cy3 labeled cell, integrated intensity was measured across a region of interest (ROI) corresponding to the cell periphery. The CTCF value was then generated by subtracting the background signal from an area equivalent to ROI from that integrated intensity.

### Purification of CERT START domain

The START domain of CERT (START_CERT_) was cloned into a pGEX-4-T1 vector, for expression in fusion at the C-terminus of a GST domain followed by a thrombin cleavage site. START_CERT_ was expressed in *E.coli* BL21-Gold(DE3) (Agilent) for 16 hr in LB-Lennox supplemented with 0.5 mM IPTG after the optical density (OD) at 600 nm reached 0.9. Pellets were obtained after a first centrifugation of the culture (30 min, 3500 g, 4°C), and second centrifugation (30 min, 3500 g, 4°C) after resuspension in cold PBS (1:10 initial culture volume). The pellet was frozen at −20 °C until subsequent protein extraction and purification. All purification steps were performed in cold and degassed TN buffer (50 mM Tris-base, 300 mM NaCl, pH 8.0 at 4 °C adjusted with HCL). From 1 L of culture, frozen bacterial pellets were resuspended in 100 mL TN buffer supplemented with 1 mM DTT (TND buffer), 2 tablets of Complete® EDTA-free protease inhibitor cocktail (Roche), 10 μM bestatin, 1 μg/mL pepstatin A, and 10 μM phosphoramidon. Cells were lysed by passing them twice through a homogenizer, EmulsiFlex-C3 (Avestin), the lysate was then doped with 200 mM PMSF, then 5 mM MgCl2 and 20 µg/mL DNAse I (Roche) and clarified by ultracentrifugation (186,000 g, 1h, 6°C), soluble protein containing supernatant was applied to Glutathione Sepharose slurry (3% v/v) and incubated 3.5 hr at 4 °C. The resin was then packed and washed 4 times with 10 volumes of TND buffer using an Econo-Pac chromatography column. Bound proteins were eluted with TND buffer supplemented with 10 mM glutathione reduced (Sigma-Aldrich). The eluate was collected, and purity determined by SDS-PAGE analysis stained with InstantBlue (Abcam). GST-CERT containing fractions were diluted to 15 mL with TN buffer and concentrated to 2 mL twice using an ultrafiltration device (MWCO 10 kDa, Amicon Ultra-15). The sample was then buffer exchanged with a PD-10 column (Cytiva) equilibrated with a fresh TN buffer. Samples were concentrated, supplemented with 10% glycerol (v/v), flash frozen in liquid N_2_ and stored at −80°C until further use. Sample purity was confirmed by mass identification on an Agilent 6230 time-of-flight mass spectrometer (TOFMS) with JetStream electrospray ionization source ESI (LC-ESI-TOFMS) after dialysis (MWCO 3.5 kDa, Slide-A-Lyzer) in low salt TN buffer (10 mM Tris-HCl, 50 mM NaCl, pH 7.4).

### *In vitro* activity of CERT

DOPC (1,2-dioleoyl-sn-glycero-3-phosphocholine), and Rhod-PE (1,2-dipalmitoyl-sn-glycero-3-phosphoethanolamine-N-(lissamine rhodamine B sulfonyl)), which were used from stock solutions in CHCl_3_ (Avanti Polar Lipids). The concentrations of NBD-dihydroceramide (N-[(E,2S,3R)- 1,3-dihydroxyoctadec-4-en-2-yl]- 12-[(4-nitro-2,1,3-benzoxadiazol-7-yl) amino]dodecanamide and NBD-deoxydihydroceramide (N-[(2S,3R,4E)-3-hydroxyoctadec-4-en-2-yl]-12-[(7-nitro-2,1,3-benzoxadiazol-4-yl)amino]dodecanamide) in methanol were determined by absorbance at 463 nm (ε = 22,000 M^-1^cm^-1^). Liposomes were prepared as previously described^91^. Briefly, desired molar ratios of lipids were combined from stock solutions in a pear-shaped flask and dried under vacuum using a rotary evaporator with a water bath set at 40 °C. The resulting lipid films were resuspended in preheated HK buffer (40 °C; 50 mM HEPES, 120 mM potassium acetate, pH 7.4, adjusted with KOH) using four sterile glass beads and vigorous vortexing for 5 minutes to ensure complete resuspension of the waxy lipids. Vesicle lamellarity was then reduced by five freeze–thaw cycles. Large unilamellar vesicles (LUVs) were subsequently obtained by extrusion through a 0.2 µm perforated polycarbonate membrane using a mini-extruder and syringes pre-heated to 60 °C. The NBD-ceramide transfer from liposome donor (L_A_) to liposomes acceptor (L_B_) was measured by recording the dequenching of NBD fluorescence on Cary Eclipse Fluorescence Spectrophotometer (Agilent). In a 500 µL rectangular quartz suprasil cuvette (Perkin Elmer), stirred with a magnetic bar, 200 µM of L_A_ liposome were added (186 µM DOPC, 10 µM C12-NBD-DHCer or C12-NBD-deoxyDHCer, 4 µM Rhodamine PE). The NBD fluorophore was recorded at 538 nm (excitation slit = 5 nm) under excitation at 473 nm (slit = 2.5 nm) with a photomultiplier set to 800 V, and measurement interval of 0.5 sec. At t = 1 min, 200 µM of L_B_ liposomes (200 µM DOPC) were added to the cuvette, a slow (∼0.23 nM / minutes) spontaneous transfer is measured over two minutes. CERT was then added at a final concentration of 200 nM and the kinetics of transfer was recorded over 12 minutes. For inhibition, CERT protein samples were incubated with five molar excesses of HPA-12 in DMSO (10 mM stock) or with 0.036% DMSO and incubated at room temperature for 5 minutes before measuring the activity. To determine the amount of lipid transported, the maximal fluorescence (Fmax) under equilibrium condition was measured by mixing 200 µM L_A-eq_ liposomes (191 µM DOPC, 5 µM C12-NBD-DHCer or C12-NBD-deoxyDHCer, 4 µM Rhodamine PE) and 200 µM L_B-eq_ liposomes (195 µM DOPC, 5 µM C12-NBD-DHCer or C12-NBD-deoxyDHCer, 4 µM Rhodamine PE). The concentration of NBD-lipid transported was calculated from the recorded fluorescence signal (F) using the following equation: [Conc] = (F - F_0_) / (F_max_ - F0) * 5; where F_0_ represents the average fluorescence over 6 seconds prior CERT injection in each independent measurement, and F_max_ corresponded to the average fluorescence over 5 minutes under equilibrium conditions after 9 minutes incubation, from three independent experiments. The initial rate was determined from the slope of a linear regression over the 4 first seconds following protein injection.

### Generation of cell lines

Stable cell lines expressing SPTLC-1^WT^ (SPT^WT^) and SPTLC-1^C133W^ (SPT^C133W^) were generated using lentiviral transduction as previously described^31^. In brief, 6 μL or lentivirus particles harboring each ORF were added to RPE-1 in 0.5 mL medium containing 6 μg/mL polybrene for 4 h before addition of 2 mL virus-free growth medium. After 24 h, the medium was changed to standard growth medium containing 5 μg/mL puromycin for 10 days, refreshing every 48 h. After puromycin selection, immunoblotting was carried out to confirm the expression of SPTLC-1 in response to doxycycline.

### Western blotting

To confirm the integration of SPTLC-1^WT^ and SPTLC-1^C133W^ alleles into hTERT-RPE-1 cell line under titratable *tet* promoter, puromycin selected cells were plated on T-25 flasks in growth medium containing 10% dialyzed FBS and doxycycline. After 48 h of induction with doxycycline at different concentrations, cells were washed twice with cold PBS and lysed in ice-cold Pierce^TM^ RIPA buffer (Thermo Fisher Scientific) supplemented with 1X Halt protease inhibitor cocktail (Thermo Fisher Scientific) and incubated for 5 min at 4°C. Cell lysates were then collected and centrifuged at 14,000 g for 15 min at 4°C. Supernatants were used to quantify protein concentration using Pierce^TM^ BCA protein assay kit (Thermo Fisher Scientific). Samples were incubated for 5 min at 95°C in 1X Laemmli buffer. 12 μg of total protein was separated on a 4-20% SDS-PAGE gel (mini-PROTEAN TGX gels; Bio-Rad) along with Precision Plus Protein Dual Color standards (Bio-Rad), and proteins were transferred onto a 0.2 µm PVDF membrane (Bio-Rad). The membrane was blocked with 5% nonfat milk in TBS buffer with 0.1% Tween-20 (TBST) for 1 h at room temperature and immunoblotted with primary antibody at 4°C overnight diluted in 5% nonfat milk, anti-SPTLC1 (Proteintech 15376-1-AP, 1:500 dilution), and anti-β-actin (Cell Signaling Technology 8H10D10, 1:1,000 dilution). The immunoblots were then washed 3 times with TBST and incubated with secondary antibody for 3 hours at room temperature (1:2,000 dilution, anti-rabbit HRP conjugate for SPTLC1; 1:2,000 dilution, anti-mouse HRP conjugate for β-actin). Specific signal was detected using SuperSignal West Pico Chemiluminescent Substrate (Thermo Fisher Scientific) and imaged with a Bio-Rad ChemiDoc XRS+ imaging station.

For analysis of ER stress response, SPT^WT^ and SPT^C133W^, cell lines were seeded in 6-well plates with a starting seeding density of 200,000 cells/well. Cells were induced with 1 μg/ml of doxycycline after 24 hours. Cells were harvested 48 hours post-induction. Protein levels were detected using the following primary antibodies: rabbit anti-IRE1-phosphorylated (Invitrogen PA1-16927, at 1:1,000), rabbit anti-AKT-phosphorylated (Invitrogen 44-621G, at 1:1,000 dilution), mouse anti-Derlin-1 (Sigma-Aldrich SAB4200148, at 1:2,000 dilution), rabbit anti-Derlin-3 (Thermo Fisher Scientific PA5-107110, at 1:2,000 dilution).

### Physiological assays

For analysis of cell line growth, RPE-1, SPT^WT^ and SPT^C133W^ lines were seeded in 96-well plates with starting density of 500 cells/well. Each cell line was seeded in 20 wells; in 10 of these wells, cells were treated with 1 μg/ml of doxycycline at time of seeding, and the other 10 wells were left untreated. Cell growth was performed by live cell imaging with a Incucyte Sx5 automated confluency imager (Satorius). Images were acquired every hour for 6 days and analyzed to determine cell confluency relative to the initial state at 0 hours.

For analysis of respiration in stable cell lines expressing SPT^WT^ and SPT^C133W^, 20,000 cells were seeded in biological replicates (N=3) into 96 well Seahorse cell culture microplates (Agilent Technologies) pretreated with fibronectin. Upon attachment, cells were induced with 1 μg/mL doxycycline for 48h prior to analysis on the Agilent Seahorse XF pro (Agilent Technologies). Subsequently, samples were analyzed using the Seahorse Cell Mito Stress Test with sequential addition of 1.5 μM oligomycin, 1 μM carbonyl cyanide-p-trifluoromethoxy phenylhydrazone (FCCP) and 0.5 μM of Rotenone/Antimycin A mixture. After the assay, cells were trypsinized and mixed with trypan blue to count the live cells for normalization of respiration rates.

### Lipid mass spectrometry

Cells were spiked with the following internal standards: 20 pmol sphinganine-d7 (Avanti Polar Lipids, Cat# 860658), deoxysphinganine-d3 (Avanti Polar Lipids, Cat# 860474), 100 pmol d18:0-d7/13:0 dihydroceramide (Avanti Polar Lipids, Cat# 330726), 200 pmol d18:1-d7/15:0 ceramide (Avanti Polar Lipids, Cat# 860681), 100 pmol d18:1-d7/15:0 glucosylceramide (Avanti Polar Lipids, Cat# 330729), 100 pmol d18:1-d7/15:0 lactosylceramide (Avanti Polar Lipids, Cat# 330727), 200 pmol sphingosine-d7 (Avanti Polar Lipids, Cat# 860657), and 200 pmol d18:1/18:1-d9 sphingomyelin (Avanti Polar Lipids, Cat# 791649) or 200 pmol sphingomyelin (d18:1/18:1)-d9 (Avanti Polar Lipids, Cat#860740). Cells were scraped in a solution containing 0.5 mL methanol and 0.5 mL water. A 100 µL aliquot of the homogenate was set aside to determine protein concentration using a BCA protein assay (Thermo Fisher Scientific). The remaining homogenate was transferred to a new Eppendorf tube, followed by the addition of 1 mL of chloroform. The samples were vortexed for 5 minutes and centrifuged at 15,000 g for 5 minutes at 4°C. The organic phase was collected, and 2 µL of formic acid was added to the polar phase, which was re-extracted with an additional 1 mL of chloroform. The organic phases were combined and dried under nitrogen.

Sphingolipids were quantified using an Agilent 6460 QQQ LC-MS/MS system. Separation was achieved on a C8 column (Spectra 3 μm C8SR 150 × 3 mm, Peeke Scientific). Dried extracts were resuspended in 100 µL of Buffer B (methanol with 1 mM ammonium formate and 0.2% formic acid), sonicated for 10 minutes, centrifuged at 15,000g for 10 minutes at 4°C, and 80 µL of the supernatant was transferred into vials for analysis. A 5 µL injection was made into the system. The mobile phase consisted of HPLC-grade water (phase A) with 2 mM ammonium formate and 0.2% formic acid, and methanol (phase B) with 1 mM ammonium formate and 0.2% formic acid. The flow rate was set at 0.5 mL/min and the gradient elution program was as follow: 82% B from 0 to 4 min, raised to 90% B over 14 min, raised to 99% B over 7 min, kept at 99% B for 2 min, and decreased to 82% B over 3 min, for a total run time of 30 min. A post-run of 10 min followed each sample allowing for the column re-equilibration. Sphingolipid species were detected using multiple reaction monitoring (MRM) of the transition from precursor to product ions^92^ with optimized collision energies and fragmentor voltages. Specific MRMs for sphinganine (SA), sphingosine (SO), and sphingosine-1-phosphate (S1P) were recorded from 0 to 10 minutes, while MRMs for dihydroceramide (DHCer), ceramide (Cer), hexosyl-ceramide (HexosylCer), lactosyl-ceramide (LactosylCer), and sphingomyelin (SM) were recorded from 10 to 30 minutes. Relative sphingolipid abundances were calculated by normalizing abundances to internal standards specific to their respective class and to protein content.

### Membrane fluidity analysis

For Laurdan experiments, cells were seeded and either induced with 1 μg/mL doxycycline for 48 h (SPT^WT^ and SPT^C133W^ cell lines) or treated with different inhibitors, prior to the staining protocol. Upon completion of treatment, cells were washed with HBSS (Gibco) and stained with 5 μM Laurdan dye (Thermo D250) for 30 min in serum-free DMEM/F12 media (Gibco). Staining solution was then replaced with complete growth media before imaging. To calculate GP values of specific secretory membranes, cells were also transfected with different organelle markers, Sec61β-mCherry for ER and SiT-mApple for Golgi apparatus or stained with CellMask Deep Red for plasma membrane.

To quantify the GP value of specific secretory membranes, cell images were acquired in 3 different spectral channels. A binary mask was first created with the image of a particular organelle marker to define the region of interest (ROI) and the resulting mask was then applied to images acquired on the ordered channel (436±18 nm) and disordered channel (498±18 nm). The GP was calculated at each pixel as described previously^51^ by utilizing the following equation:

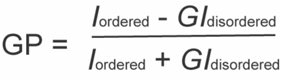

In this equation, *I* represents the intensity of each pixel in the image obtained from the specified spectral channel. The G factor compensates for experimental variations and ensures that GP measurements are comparable across different experiments. The G factor was calculated as previously described^51^ using diluted dye in pure DMSO. Visual heatmaps were generated with Python’s seaborn library to show the spatial distribution of GP values across the organelle.

### ERES size quantification

Each endoplasmic reticulum exit site (ERES) was cropped from the original image. Approximately the same number of exit sites were cropped from each image. Only ones with no overlap with other sites were chosen. The length of each square pixel was defined from the dimensions of the image given by ImageJ with the chosen microscope setting. To define the number of pixels comprising each ERES, the region of interest was selected by thresholding. Otsu’s threshold was used to calculate an appropriate threshold value for the image. Area of each ERES was measured by converting from the number of pixels to square microns.

### Synchronized cargo release

For live-cell experiments, cells were transiently transfected with single-plasmid RUSH systems containing both hook and cargo. Cargoes were released upon addition of 40 μM of biotin on the microscope stage. Time series were acquired with 5- or 10-minute intervals. Signal associated with the Golgi region was analyzed after acquisition by integrating signal intensity in a hand-drawn region corresponding to the apparent Golgi region observed during the time-series. For this, Fiji was used alongside the StackReg plugin. This integrated intensity was normalized to the total cargo intensity across the cell.

For fixed-cell experiments, cells were fixed at discrete time points with cold 4% paraformaldehyde and 4% sucrose for 20 min at room temperature in the dark. Fixed cells were then washed with PBS followed by permeabilization and blocking in 0.05% saponin and 1% BSA in PBS. Permeabilized cells were then incubated with anti-GM130 primary antibody (Abcam Ab52649, 1:500 dilution) in the same buffer at 4°C overnight. After three washes, the samples were incubated with a AlexaFluor647-labeled secondary antibody (Abcam Ab150091, 1:500 dilution) for 1 hr. The stained samples were washed three times and imaged at room temperature. To quantify the fraction of cargo in the Golgi compartment at a given time point, images of the Golgi marker (GM130) and the fluorescent cargo were segmented using Otsu’s method. The fraction of fluorescent cargo signal present in the Golgi was then calculated by dividing the sum of fluorescent cargo pixel intensities overlapping with the Golgi marker by the total fluorescent cargo pixel intensity.

## Supporting Information

**Figure S1.**
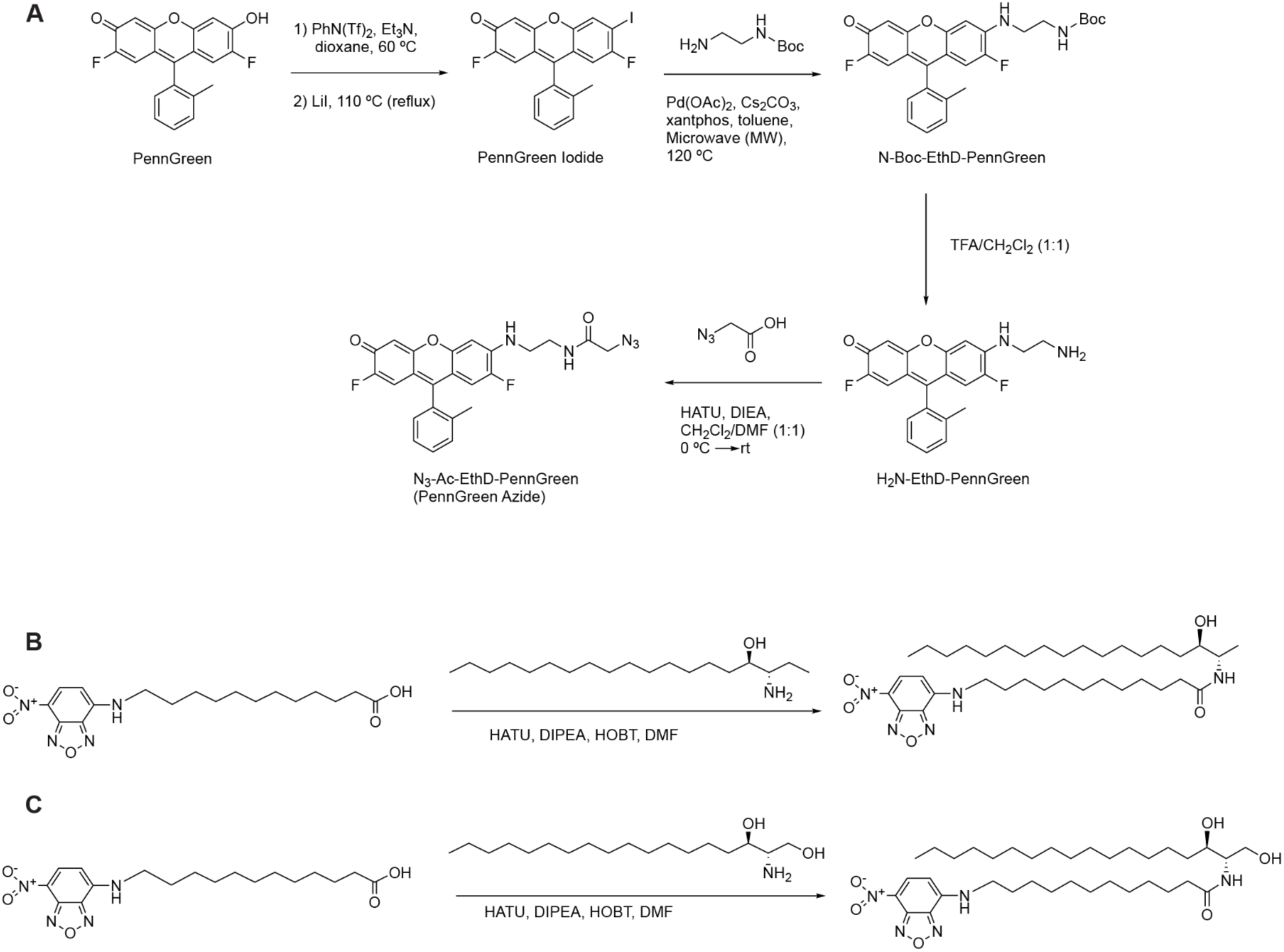
Synthesis of new compounds used in this study. PennGreen-azide for early secretory membrane labeling. **A.** Synthesis of the PennGreen-azide probe used for early secretory membrane labeling in this study. **B.** Synthesis of C12-NBD-deoxyDHCer used for CERT lipid transfer assay. **C.** Synthesis of C12-NBD-DHCer. Details provided in the Supplementary Information.

**Figure S2.**
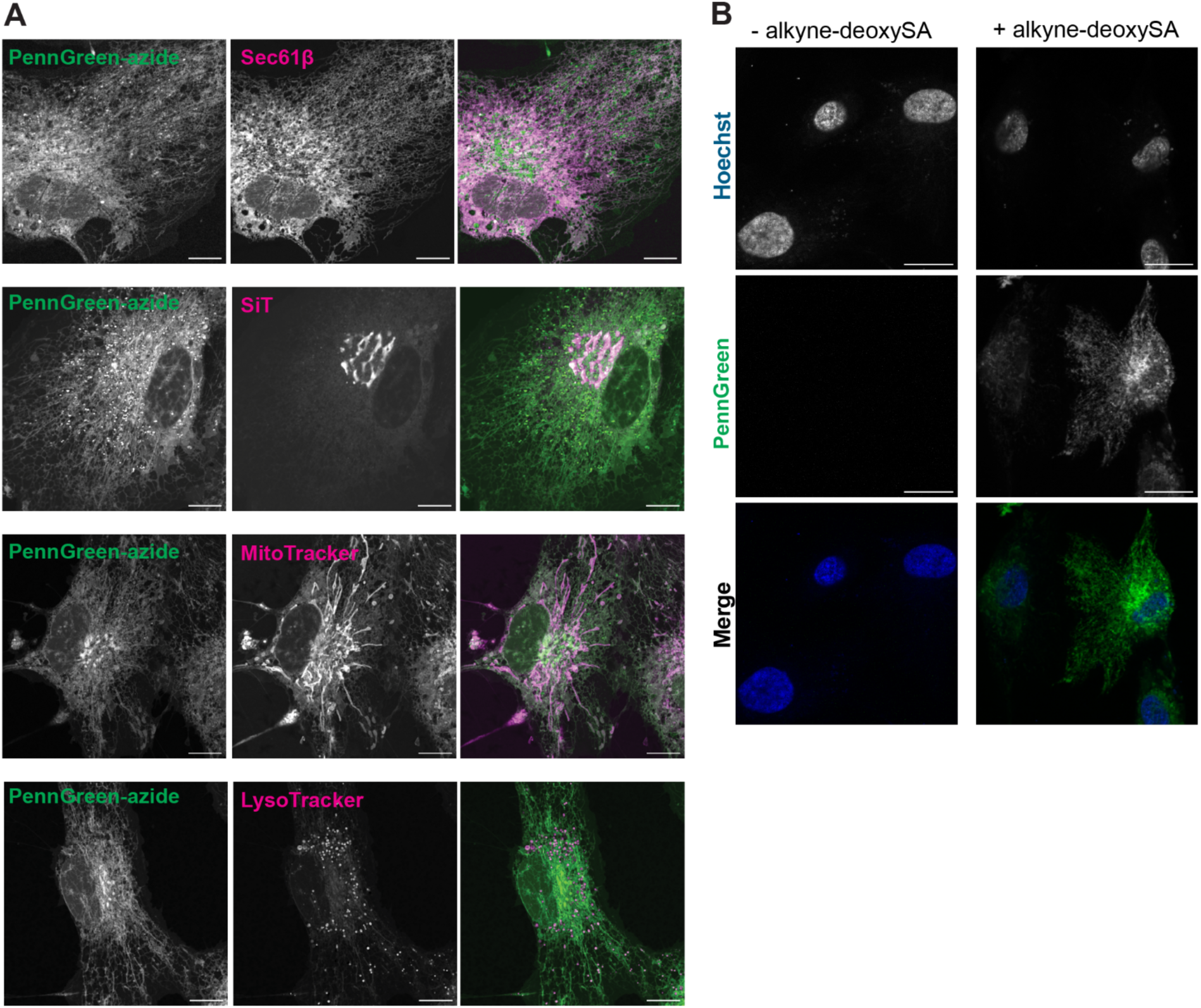
PennGreen-azide validation. **A.** Localization of PennGreen-azide in live cells. RPE-1 cells were transfected with Sec61β (ER marker), SiT (Golgi marker) or stained with MitoTracker (mitochondrial marker), Lysotracker (lysosome marker) prior to the treatment with PennGreen-azide for 45 min. Scale bars, 10 μm. **B.** PennGreen staining is dependent on the alkyne substrate. RPE-1 fed without (left) or with (right) 0.1 μM alkyne-deoxySA were prepared and imaged identically. Only cells fed the alkyne substrate show PennGreen signal after CuAAC reaction and washing steps. Scale bars, 20 μm.

**Figure S3.**
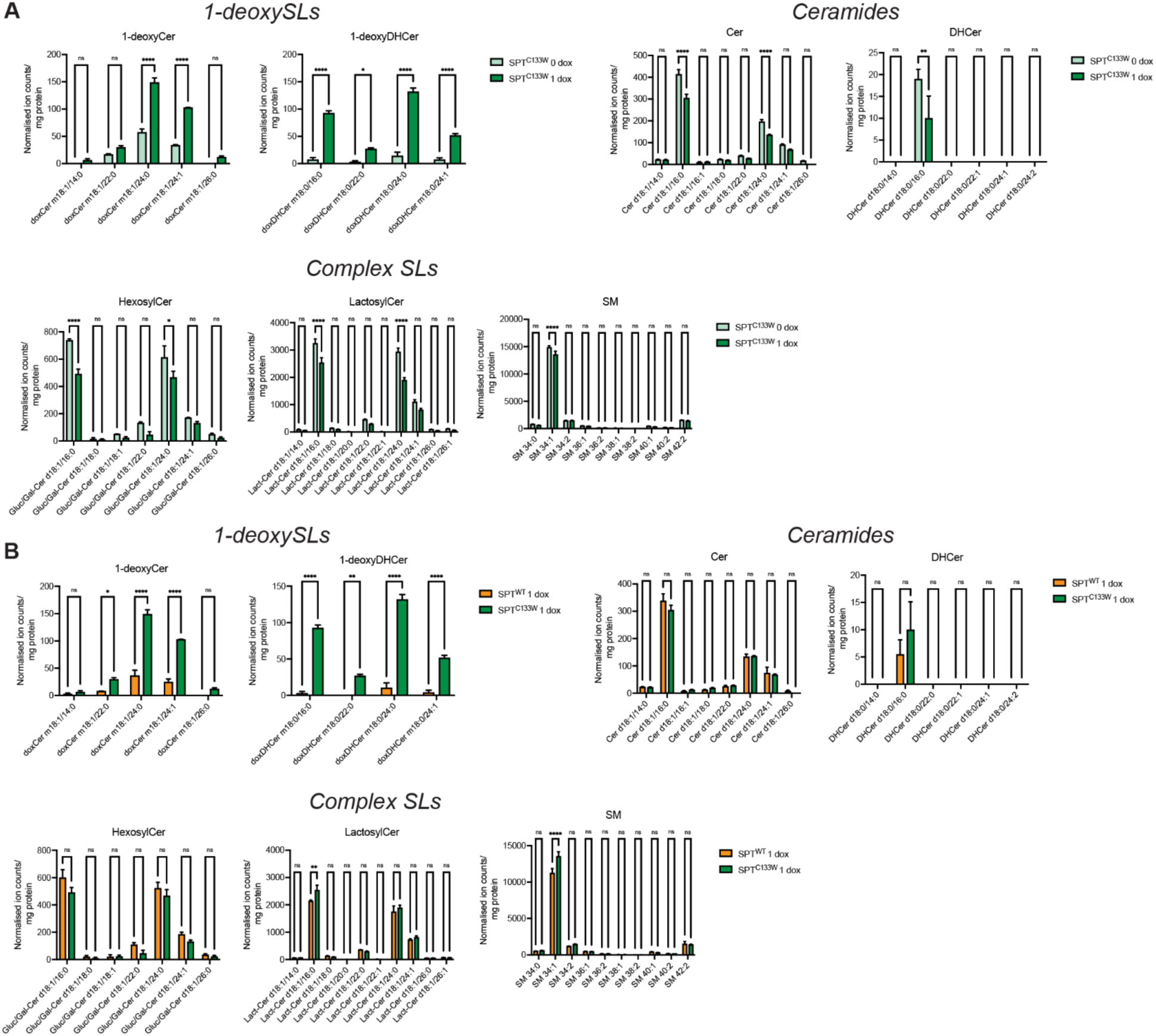
Species-level breakdown of SL changes in SPT^C133W^ cells. **A.** Comparison between uninduced and maximally induced (1 μg/mL, 48 h) SPT^C133W^. **B.** Comparison between induced (1 μg/mL, 48 h) SPT^WT^ and SPT^C133W^. Error bars indicate SEM (extracts from N = 3 independent culture dishes). *, P < 0.05; ** P < 0.01; ***, P < 0.001; ****, P < 0.0001; ns, no significance by 2-way ANOVA.

**Figure S4.**
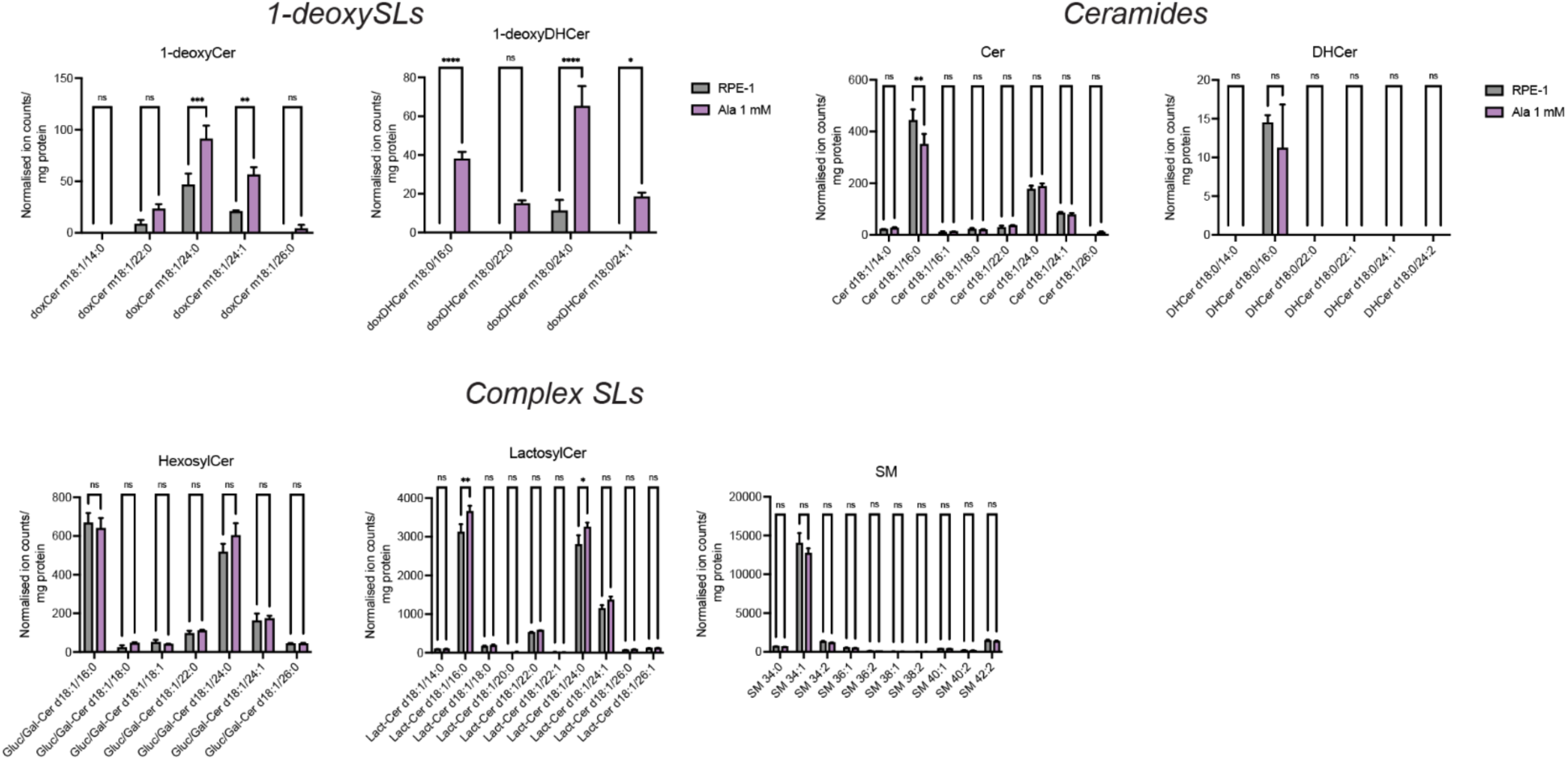
Species-level breakdown in Ala-supplemented cells. Comparison between Ala (1 mM, 48 h) supplemented and a DMSO control RPE-1. Error bars indicate SEM (extracts from N = 3 independent culture dishes). *, P < 0.05; ** P < 0.01; ***, P < 0.001; ****, P < 0.0001; ns, no significance by 2-way ANOVA.

**Figure S5.**
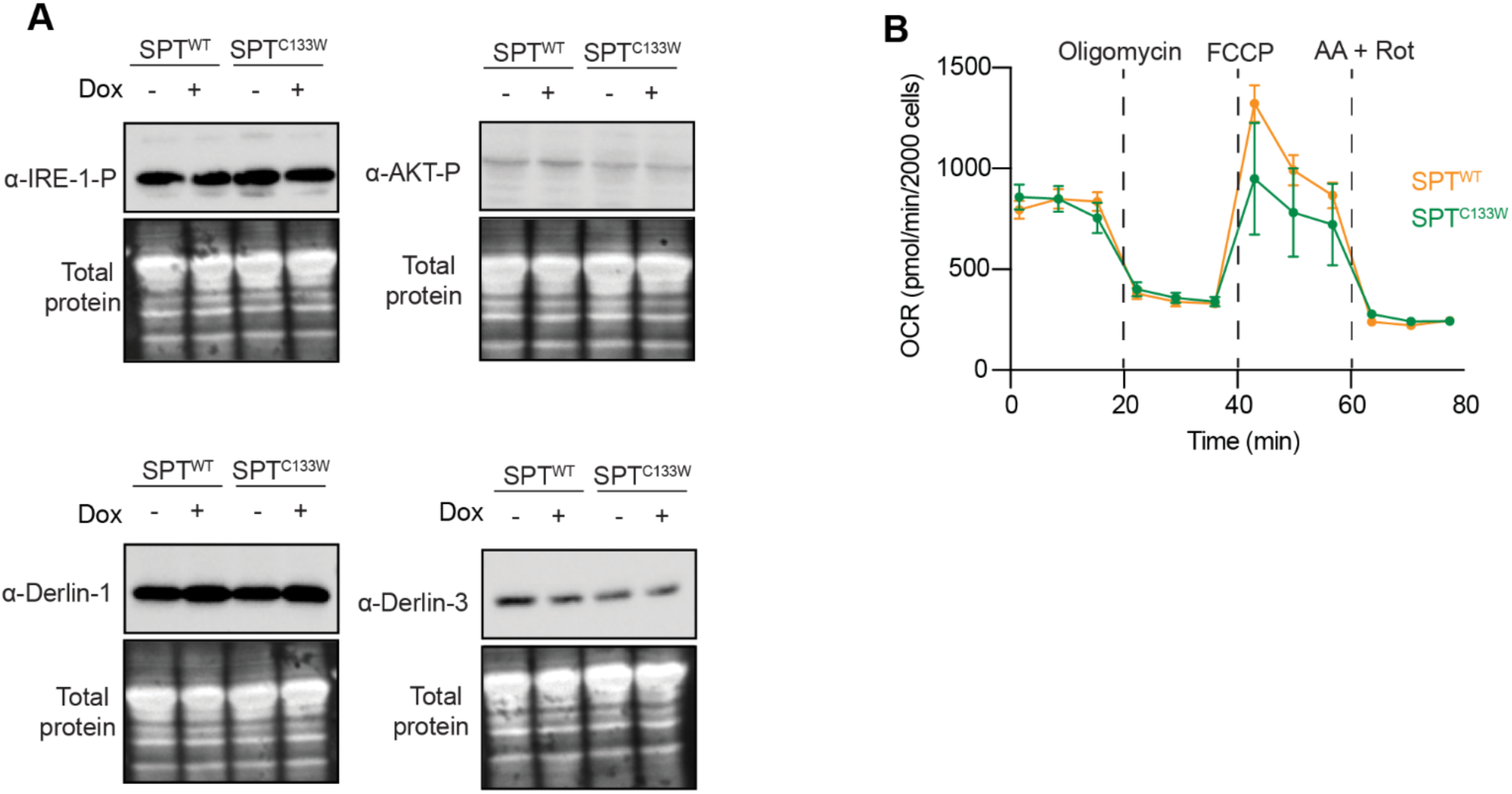
SPT^C133W^ cells do not display classical markers for ER stress or mitochondrial dysfunction. **A.** Western blots for levels of UPR and ERAD markers in uninduced or induced (1 μg/mL, 48 h) SPT^WT^ and SPT^C133W^ cells. No major differences are observed as a function of either induction or the SPTLC-1 variant expression. **B.** Seahorse respirometry of induced (1 μg/mL, 48 h) SPT^WT^ and SPT^C133W^ cells showing levels of oxygen consumption for basal respiration (0-20 min), proton leak (20-40 min), uncoupled respiration (40-60 min), and background oxygen consumption (60-80 min). Error bars indicate SEM for N=3 independent cultures. No significant differences are observed.

**Figure S6.**
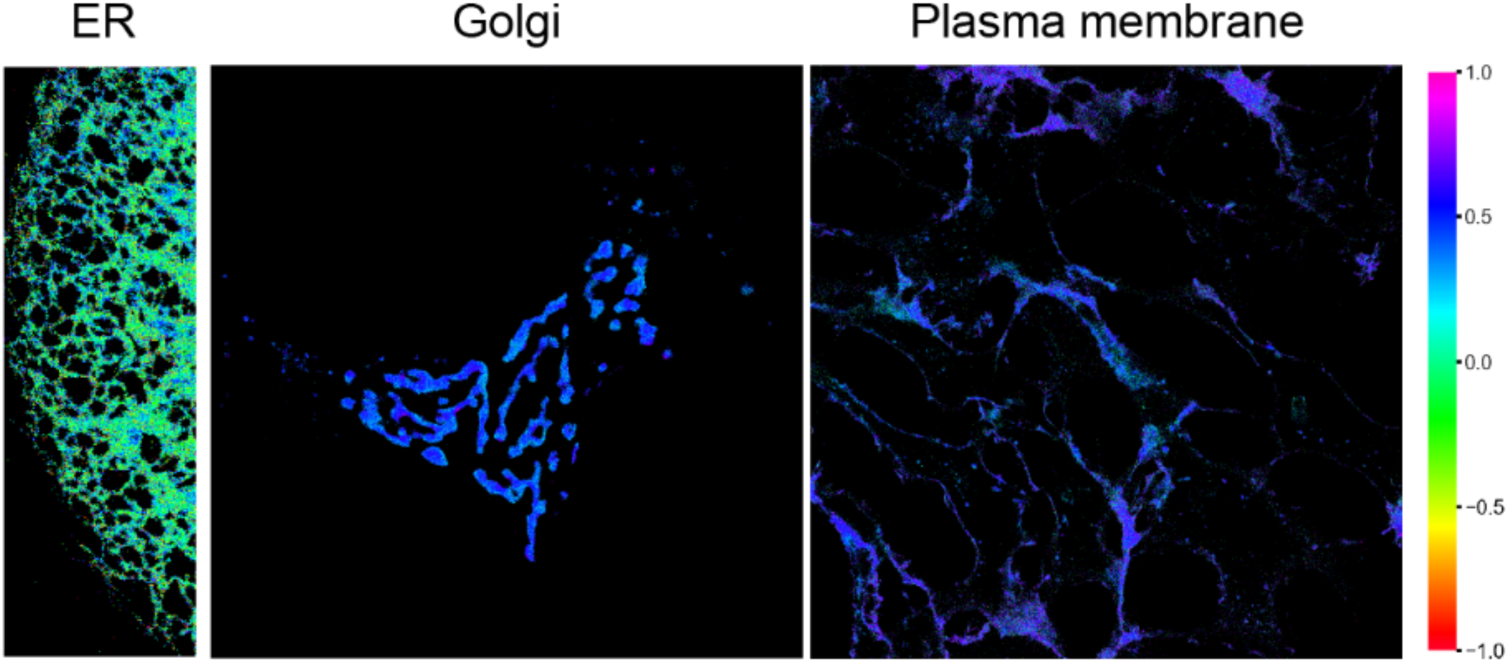
Measuring ordering in the ER, Golgi, and the Plasma Membrane using Laurdan GP heatmaps. Example heatmaps of RPE-1 cells are shown obtained from Laurdan imaging of secretory organelles by following the pipeline indicated on Figure 3. Markers are masked using Sec61β-mCherry (ER), SiT-mApple (Golgi), or CellMask Deep Red (PM). The GP images are false colored according to each pixel’s GP value with the range indicated by the color scale to the right.

**Figure S7.**
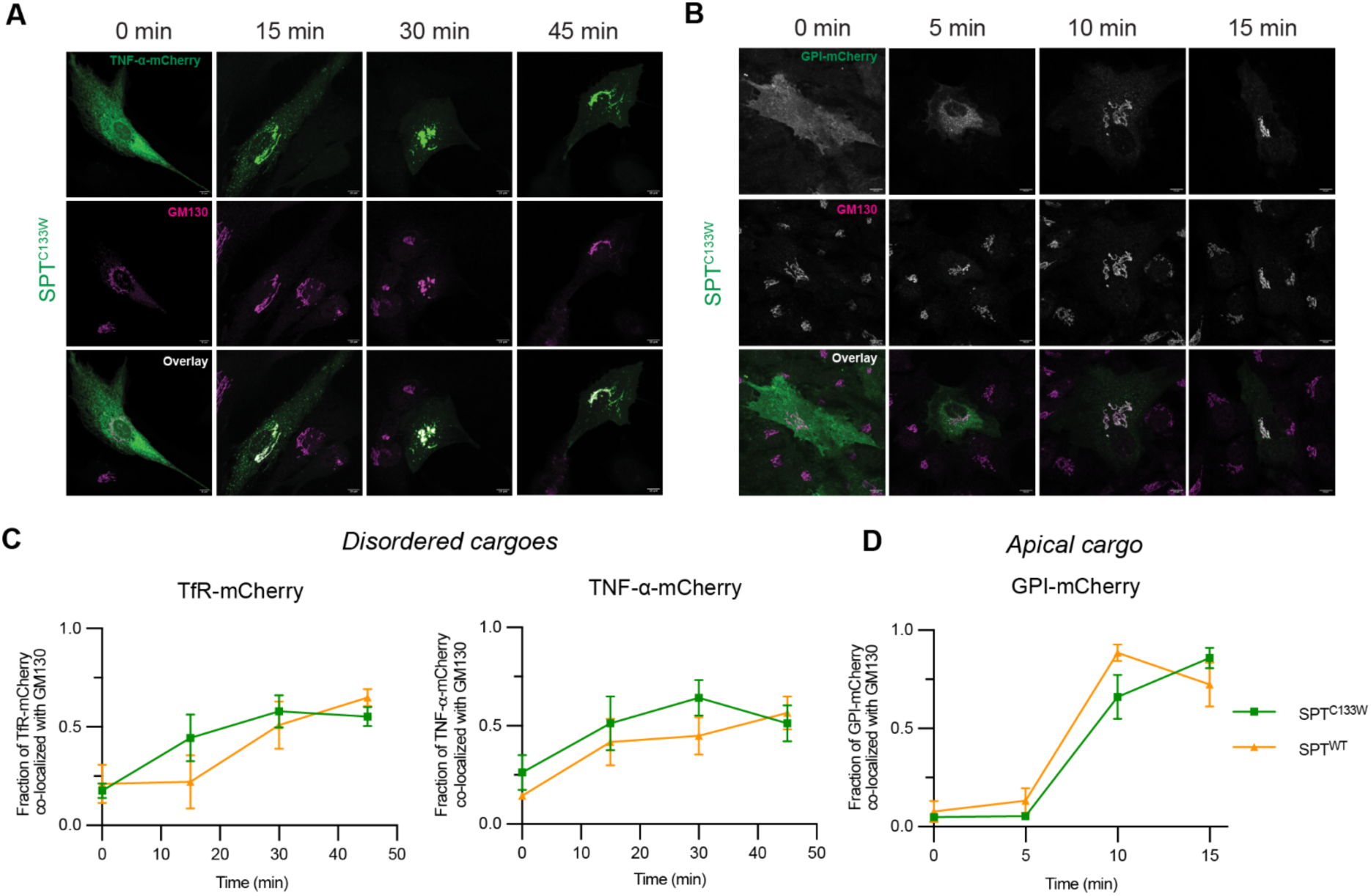
RUSH experiments in fixed cells show cargo-dependent kinetic effects of SPT^C133W^ expression. **A.** Example of RUSH experiment with TNF-α-mCherry as the cargo in SPT^C133W^ cells. The cargo starts distributed across the ER (0 minutes). Co-localization to the Golgi marker GM130 is apparent at 15 minutes and peaks at 30 minutes before decreasing due to exit from the Golgi at 45 min. Scale bars, 10 μm. **B.** Example of RUSH experiment with GPI-mCherry as the cargo in SPT^C133W^ cells. The cargo starts distributed across the ER (0 minutes), then segregates to punctae within the ER (ERES) at 5 minutes. Co-localization to the Golgi marker GM130 is apparent at 10 minutes and peaks at 15 minutes. Scale bars, 10 μm. **C.** Quantification of disordered cargoes (left: TfR-mCherry, right: TNF-α-mCherry) with anti-GM130 in induced (1 μg/mL dox, 48 h) SPT^WT^ and SPT^C133W^ cells. In both cases, SPT^C133W^ cells show faster kinetics. Each time point for each condition and cargo represents cells from a single RUSH transfected dish. Error bars show 95% CI across cells. Number of cells for TfR-mCherry - SPT^WT^: 0 min, 23; 15 min, 24; 30 min, 34; 45 min, 32. TfR-mCherry - SPT^C133W^: 0 min, 28; 15 min, 33; 30 min, 35; 45 min, 44. TNF-α-mCherry - SPT^WT^: 0 min, 31; 15 min, 31; 30 min, 30; 45 min, 26. TNF-α-mCherry - SPT^C133W^: 0 min, 32; 15 min, 28; 30 min, 31; 45 min, 30. **D.** Quantification of an apical cargo (GPI-mCherry), where SPT^C133W^ cells show slower ER release kinetics. Each time point for each condition and cargo represents cells from a single RUSH transfected dish. Error bars show 95% CI across cells. Number of cells for GPI-mCherry - SPT^WT^: 0 min, 29; 5 min, 40; 10 min, 40; 15 min, 32. GPI-mCherry - SPT^C133W^: 0 min, 26; 5 min, 41; 10 min, 42; 15 min, 27.

## Notes

### Competing Interest Statement

The authors have declared no competing interest.

### Summary of Updates

Corrections made in the methods section for the antibodies used; Figure 1 updates to correct offset errors; abstract text corrected to match manuscript file.

